# Spotless: a reproducible pipeline for benchmarking cell type deconvolution in spatial transcriptomics

**DOI:** 10.1101/2023.03.22.533802

**Authors:** Chananchida Sang-aram, Robin Browaeys, Ruth Seurinck, Yvan Saeys

**Affiliations:** Data mining and Modelling for Biomedicine, VIB Center for Inflammation Research, Ghent, Belgium; Department of Applied Mathematics, Computer Science and Statistics, Ghent University, Ghent, Belgium

## Abstract

Spatial transcriptomics (ST) is an emerging field that aims to profile the transcriptome of a cell while keeping its spatial context. Although the resolution of non-targeted ST technologies has been rapidly improving in recent years, most commercial methods do not yet operate at single-cell resolution. To tackle this issue, computational methods such as deconvolution can be used to infer cell type proportions in each spot by learning cell type-specific expression profiles from reference single-cell RNA-sequencing (scRNA-seq) data. Here, we benchmarked the performance of 11 deconvolution methods using 63 silver standards, three gold standards, and two case studies on liver and melanoma tissues. The silver standards were generated using our novel simulation engine *synthspot*, where we used seven scRNA-seq datasets to create synthetic spots that followed one of nine different biological tissue patterns. The gold standards were generated using imaging-based ST technologies at single-cell resolution. We evaluated method performance based on the root-mean-squared error, area under the precision-recall curve, and Jensen-Shannon divergence. Our evaluation revealed that method performance significantly decreases in datasets with highly abundant or rare cell types. Moreover, we evaluated the stability of each method when using different reference datasets and found that having sufficient number of genes for each cell type is crucial for good performance. We conclude that while cell2location and RCTD are the top-performing methods, a simple off-the-shelf deconvolution method surprisingly outperforms almost half of the dedicated spatial deconvolution methods. Our freely available Nextflow pipeline allows users to generate synthetic data, run deconvolution methods and optionally benchmark them on their dataset (https://github.com/saeyslab/spotless-benchmark).

## Introduction

Unlike single-cell sequencing, spatial transcriptome profiling technologies can uncover the location of cells, adding another dimension to the data that is essential for studying systems biology, e.g., cell-cell interactions and tissue architecture [1]. These approaches can be categorized into imaging-or sequencing-based methods, each of them offering complementary advantages [2]. As imaging-based methods use labeled hybridization probes to target specific genes, they offer subcellular resolution and high capture sensitivity, but are limited to a few hundreds or thousands of genes [3], [4]. On the other hand, sequencing-based methods offer an unbiased and transcriptome-wide coverage by using capture oligonucleotides with a polydT sequence [5], [6]. These oligos are printed in clusters, or spots, each with a location-specific barcode that allows identification of the originating location of a transcript. While the size of these spots initially started at 100 µm in diameter, some recent sequencing technologies have managed to reduce their size to the subcellular level, thus closing the resolution gap with imaging technologies [7], [8]. However, these spots do not necessarily correspond to individual cells, and therefore, computational methods remain necessary to determine the cell-type composition of each spot.

Deconvolution and mapping are two types of cell type composition inference tools that can be used to disentangle populations from a mixed gene expression profile [9]. In conjunction with the spatial dataset, a reference scRNA-seq dataset from an atlas or matched sequencing experiment is typically required to build cell type-specific gene signatures. Deconvolution infers the proportions of cell types in a spot by utilizing a regression or probabilistic framework, and methods specifically designed for ST data often incorporate additional model parameters to account for spot-to-spot variability. On the other hand, mapping approaches score a spot for how strongly its expression profile corresponds to those of specific cell types. As such, deconvolution returns the proportion of cell types per spot, and mapping returns the probability of cell types belonging to a spot. Unless otherwise specified, we use the term “deconvolution” to refer to both deconvolution and mapping algorithms in this study.

Although there have recently been multiple benchmarking studies [10]–[12], several questions remain unanswered. First, the added value of algorithms specifically developed for the deconvolution of ST data has not been evaluated by comparing them to a baseline or bulk deconvolution method. Second, it is unclear which algorithms are better equipped to handle challenging scenarios, such as the presence of a highly abundant cell type throughout the tissue or the detection of a rare cell type in a single region of interest. Finally, the stability of these methods to variations in the reference dataset arising from changes in technology or protocols has not been assessed.

In this study, we address these gaps in knowledge and provide a comprehensive evaluation of 11 deconvolution methods in terms of performance, stability, and scalability (**Figure 1**). The tools include eight spatial deconvolution methods (cell2location [13], DestVI [14], DSTG [15], RCTD [16], SpatialDWLS [17], SPOTlight [18], stereoscope [19], and STRIDE [20]), one bulk deconvolution method (MuSiC [21]), and two mapping methods (Seurat [22] and Tangram [23]). For all methods compared, we discussed with the original authors in order to ensure that their method was run in an appropriate setting and with good parameter values. We also compared method performance with two baselines: a “null distribution” based on random proportions drawn from a Dirichlet distribution, and predictions from the non-negative least squares (NNLS) algorithm. We evaluated method performance on a total of 66 synthetic datasets (63 silver standards and three gold standards) and two application datasets. Our benchmarking pipeline is completely reproducible and accessible through Nextflow (<u>github.com/saeyslab/spotless-benchmark</u>). Furthermore, each method is implemented inside a Docker container, which enables users to run the tools without requiring prior installation.

**Figure 1.**
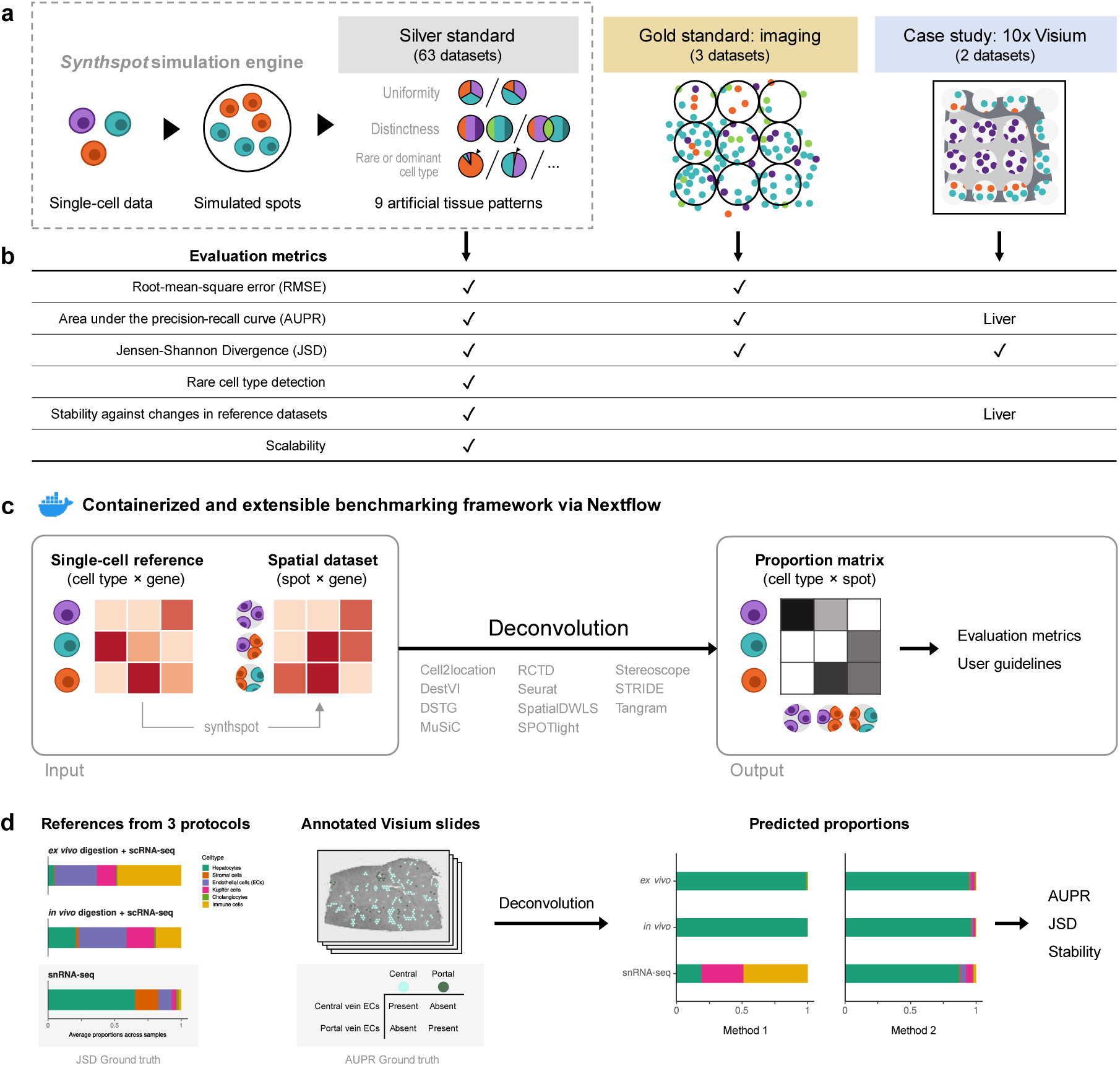
Overview of the benchmark. **(a)** The datasets used consist of silver standards generated from single-cell RNA-seq data, gold standards from imaging-based data, and two case studies on liver and melanoma. Our simulation engine *synthspot* enables the creation of artificial tissue patterns. **(b)** We evaluated deconvolution methods on three overall performance metrics (RMSE, AUPR, and JSD), and further checked specific aspects of performance, i.e., how well methods detect rare cell types and handle reference datasets from different sequencing technologies. For the case studies, the AUPR and stability are only evaluated on the liver dataset. **(c)** Our benchmarking pipeline is entirely accessible and reproducible through the use of Docker containers and Nextflow. **(d)** To evaluate performance on the liver case study, we leveraged prior knowledge of the localization and composition of cell types to calculate the AUPR and JSD. We also investigated method performance on three different sequencing protocols.

## Results

### Synthspot allows simulation of artificial tissue patterns

Synthetic spatial datasets are commonly generated by developers of deconvolution methods as part of benchmarking their algorithms against others. However, these synthetic datasets typically have spots with random compositions that do not reflect the reality of tissue regions with distinct compositions, such as layers in the brain. On the other hand, general-purpose simulators are more focused on other inference tasks, such as spatial clustering and cell-cell communication, and are usually unsuitable for deconvolution. For instance, generative models and kinetic models like those of scDesign3 [24] and scMultiSim [25] are computationally intensive and unable to model entire transcriptomes. SRTSim [26] focuses on modeling gene expression trends and does not explicitly model tissue composition, while spaSim [27] only models tissue composition without gene expression. To overcome these limitations, we developed our own simulator called synthspot that can generate synthetic tissue patterns with distinct regions, allowing for more realistic simulations (<u>github.com/saeyslab/synthspot</u>). We validated that our simulation procedure accurately models real data characteristics and that method performances are comparable between synthetic and real tissue patterns (**Supplementary Note 1**).

Within a synthetic dataset, *synthspot* creates artificial regions in which all spots belonging to the same region have the same frequency priors. Frequency priors correspond to the likelihood in which cells from a cell type will be sampled, and therefore, spots within the same region will have similar compositions. These frequency priors are influenced by the chosen artificial tissue pattern, or *abundance pattern*, which determines the uniformity, distinctness, and rarity of cell types within and across regions (**Figure 1a, Figure S1**). For example, the *uniform* characteristic will sample the same number of cells for all cell types in a spot, while *diverse* samples differing number of cells. The *distinct* characteristic constrains a cell type to only be present in one region, while *overlap* allows it to be present in multiple regions. Additionally, the *dominant cell type* characteristic randomly assigns a dominant cell type that is at least 5-15 times more abundant than others in each spot, while *rare cell type* does the opposite to create a cell type that is 5-15 times less abundant. The different characteristics can be combined in up to nine different abundance patterns, each representing a plausible biological scenario.

### Cell2location and RCTD are the top performers in synthetic data

Our synthetic spatial datasets consist of 63 silver standards generated from *synthspot* and three gold standards generated from imaging data with single-cell resolution (**Supplementary Table 1a-b**). We generated the silver standards using seven publicly available scRNA-seq datasets and nine abundance patterns. The scRNA-seq datasets consisted of four mouse brain tissues (cortex, hippocampus, and two cerebellum), mouse kidney, mouse skin cancer (melanoma), and human skin cancer (squamous cell carcinoma). Half of the cells from each scRNA-seq dataset were used to generate the synthetic spots and the other half used as the reference for deconvolution. This split was stratified by cell type. We generated 10 replicates for each of the 63 silver standards, with each replicate containing around 750 spots (**Figure S2**). For the gold standard, we used two seqFISH+ sections of mouse brain cortex and olfactory bulb (63 spots with 10,000 genes each) and one STARMap section of mouse primary visual cortex (108 spots with 996 genes) [3], [28]. We summed up counts from cells within circles of 55-µm diameter, which are the size of spots in the 10x Visium commercial platform.

We evaluated method performance with the root-mean-square error (RMSE), area under the precision-recall curve (AUPR), and Jensen-Shannon divergence (JSD) (**Supplementary Note 2**). The RMSE measures how numerically accurate the predicted proportions are, the AUPR measures how well a method can detect whether a cell type is present or absent, and the JSD measure similarities between two probability distributions.

RCTD and cell2location were the top two performers across all metrics in the silver standards, followed by SpatialDWLS, stereoscope, and MuSiC (**Figure 2b**, **Figure 3a**). All other methods ranked worse than NNLS in at least one metric. For each silver standard, method rankings were determined using the median value across 10 replicates. We observed that method performances were more consistent between abundance patterns than between datasets (**Figure S3-S5**). Most methods had worse performance in the two abundance patterns with a dominant cell type, and there was considerable performance variability between replicates due to different dominant cell types being selected in each replicate. Only RCTD and cell2location consistently outperformed NNLS in all metrics in these patterns (**Figure S6**).

**Figure 2.**
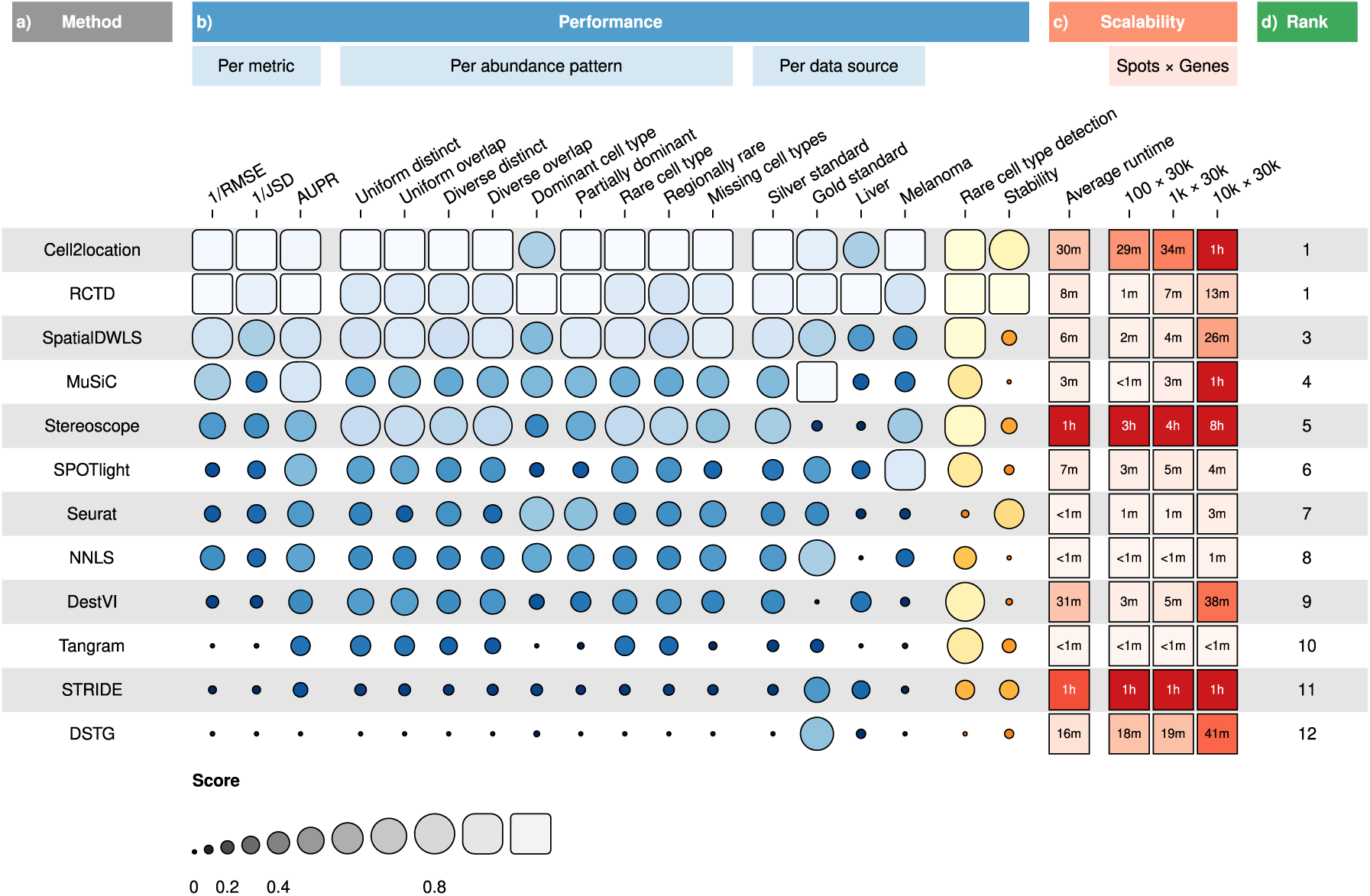
(**a**) Methods ordered according to their overall rankings **(d)**, determined by the aggregated rankings of performance and scalability. **(b)** Performance of each method across metrics, artificial abundance patterns in the silver standard, and data sources. The ability to detect rare cell types and stability against different reference datasets are also included. **(c)** Average runtime across silver standards and scalability on increasing dimensions of the spatial dataset.

**Figure 3.**
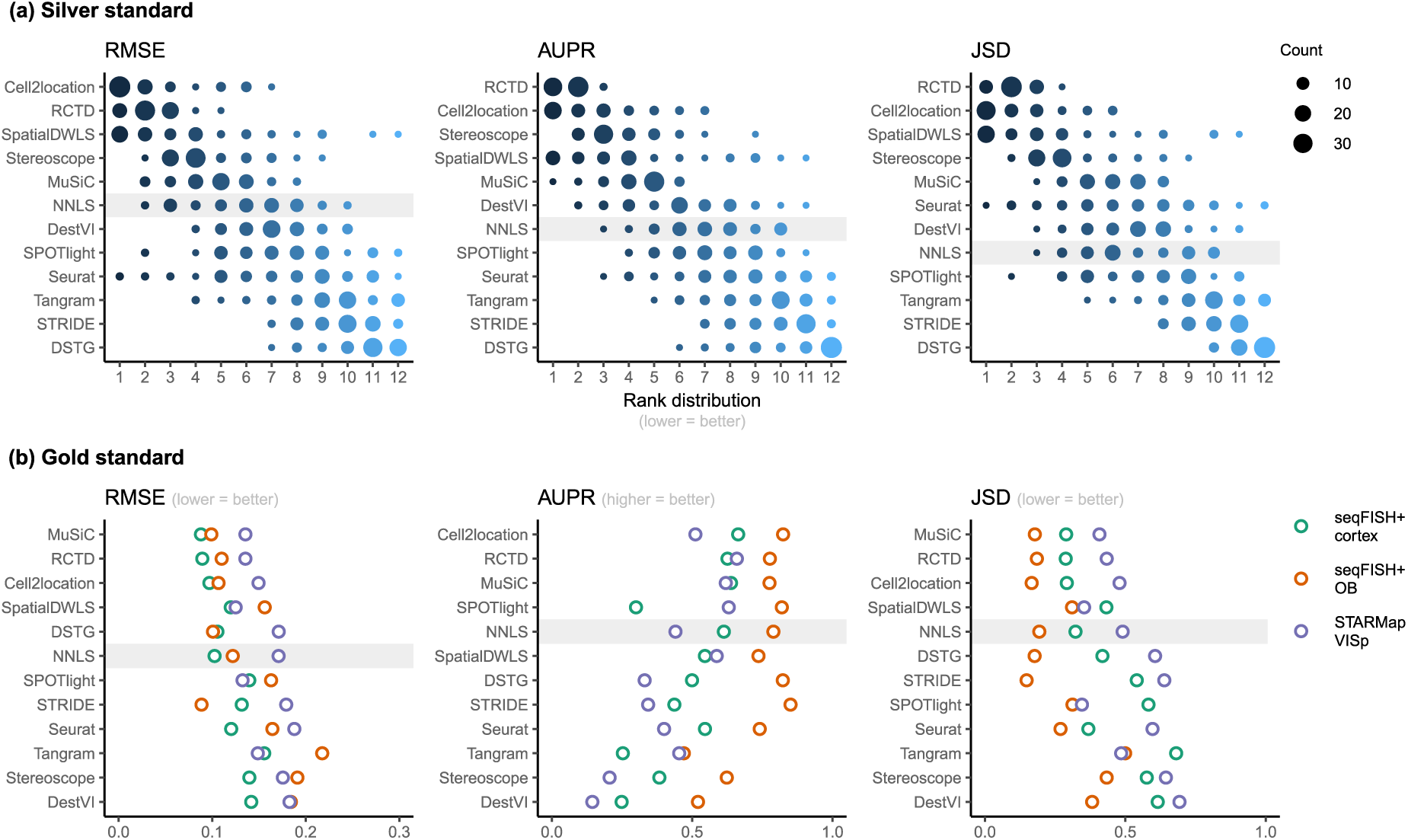
Method performance on synthetic datasets, evaluated using root-mean-squared error (RMSE), area under the precision-recall curve (AUPR), and Jensen-Shannon divergence (JSD). Non-negative least squares (NNLS) is shaded as a baseline algorithm. Methods are ordered based on the summed ranks across all 63 and three datasets, respectively. **(a)** The rank distribution of each method across all 63 silver standards, based on the best median value across ten replicates for that standard. **(b)** Gold standards of two seqFISH+ datasets and one STARMap dataset. We took the average over seven field of views for the seqFISH+ dataset.

For the gold standards, cell2location, MuSiC, and RCTD are the top three performers as well as the only methods to outrank NNLS in all three metrics (**Figure 3b**). As each seqFISH+ dataset consisted of seven field of views (FOVs), we used the average across FOVs as the representative value to be ranked for that dataset. Several FOVs were dominated by one cell type (**Figure S7**), which was similar to the dominant cell type abundance pattern in our silver standard. Consistent with the silver standard results, half of the methods did not perform well in these FOVs. In particular, SPOTlight, DestVI, stereoscope, and Tangram tended to predict less variation between cell type abundances. DSTG, SpatialDWLS, and Seurat predicted the dominant cell type in some FOVs but did not predict the remaining cell type compositions accurately. Most methods performed worse in the STARMap dataset except for SpatialDWLS, SPOTlight, and Tangram.

### Detecting rare cell types remains challenging even for top-performing methods

Lowly abundant cell types often play an important role in development or disease progression, as in the case of stem cells and progenitor cells or circulating tumor cells [29]. As the occurrence of these cell types are often used to create prognostic models of patient outcomes, the accurate detection of rare cell types is a key aspect of deconvolution tools [30], [31]. Here, we focus on the two rare cell type patterns in our silver standard (rare cell type *diverse* and *regional rare cell type diverse*), in which a rare cell type is 5-15 times less abundant than other cell types in all or one synthetic region, respectively (**Figure S1**). This resulted in 14 synthetic datasets (seven scRNA-seq datasets × two abundance patterns) with 10 replicates each. We only evaluated methods based on the AUPR of the rare cell type, using the average AUPR across the 10 replicates as the representative value for each dataset. We did not include the RMSE and JSD or consider other cell types, because in prognostic models, the presence of rare cell types is often of more importance than the magnitude of the abundance itself. Therefore, it is more relevant that methods are able to correctly rank spots with and without the rare cell type.

In line with our previous analysis, RCTD and cell2location were also the best at predicting the presence of lowly abundant cell types (**Figure 4a**). There is a strong correlation between cell type abundance and AUPR, as clearly demonstrated when plotting the precision-recall curves of cell types with varying proportions (**Figure 4b**). While most methods can detect cell types with moderate or high abundance, they usually have lower sensitivity for rare cell types, and hence lower AUPRs. Upon visual inspection of precision-recall curves at decreasing abundance levels, we found a similar pattern across all silver standards (**Figure S8**). Nonetheless, we also observe that the AUPR is substantially lower if the rare cell type is only present in one region and not across the entire tissue, indicating that prevalence is also an important factor in detection.

**Figure 4.**
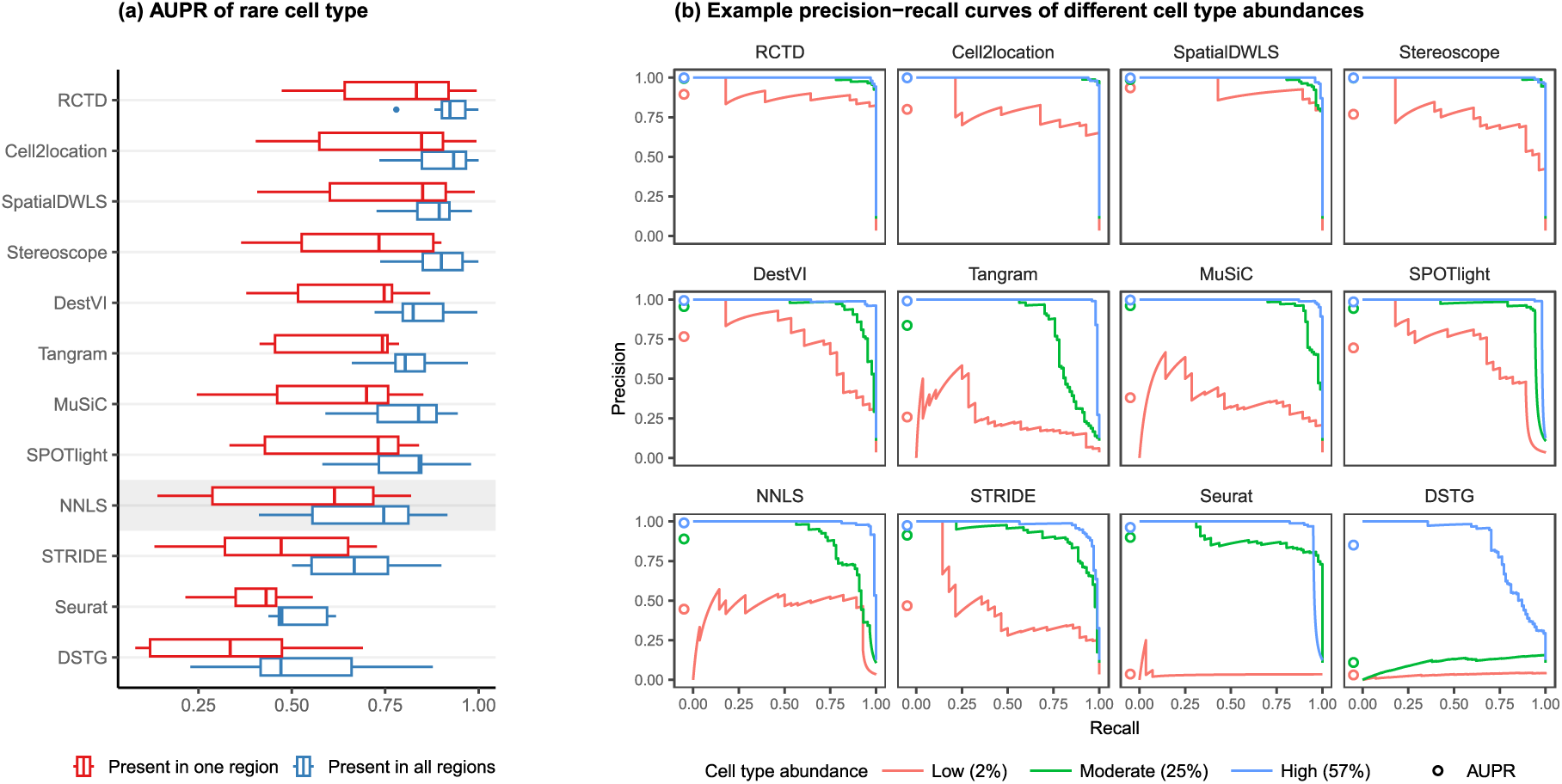
Detection of the rare cell type in the two *rare cell type* abundance patterns. **(a)** Area under the precision-recall curve (AUPR) across the seven scRNA-seq datasets, averaged over ten replicates. Methods generally have better AUPR if the rare cell type is present in all regions compared to just one region. **(b)** An example on one silver standard replicate demonstrates that most methods can detect moderately and highly abundant cells, but their performance drops for lowly abundant cells.

### Technical variability between reference scRNA-seq and spatial datasets can significantly impact predictions

Since deconvolution predictions exclusively rely on signatures learned from the scRNA-seq reference, it should come as no surprise that the choice of the reference dataset has been shown to have the largest impact on bulk deconvolution predictions [32]. Hence, deconvolution methods should ideally also account for platform effects, i.e., the capture biases that may occur from differing protocols and technologies being used to generate scRNA-seq and ST data.

To evaluate the stability of each method against reference datasets from different technological platforms, we devised the *inter*-dataset scenario, where we provided an alternative reference dataset to be used for deconvolution, in contrast to the *intra-*dataset analysis done previously, where the same reference dataset was used for both spot generation and deconvolution. We tested this on the brain cortex (SMART-seq) and two cerebellum (Drop-seq and 10x Chromium) silver standards. For the brain cortex, we used an additional 10x Chromium reference from the same atlas [33]. To measure stability, we computed the JSD between the proportions predicted from the intra-and inter-dataset scenario.

Except for MuSiC, we see that methods with better performance–cell2location, RCTD, SpatialDWLS, and stereoscope–were also more stable against changing reference datasets (**Figure 5**). Out of these four, only SpatialDWLS did not account for a platform effect in its model definition. Cell2location had the most stable predictions, ranking first in all three datasets, while both NNLS and MuSiC were consistently in the bottom three. For the rest of the methods, stability varied between datasets. DestVI performed well in the cerebellum datasets but not the brain cortex, and SPOTlight had the opposite pattern. As expected, deconvolution performance also generally worsens in the inter-dataset scenario (**Figure S10**).

**Figure 5.**
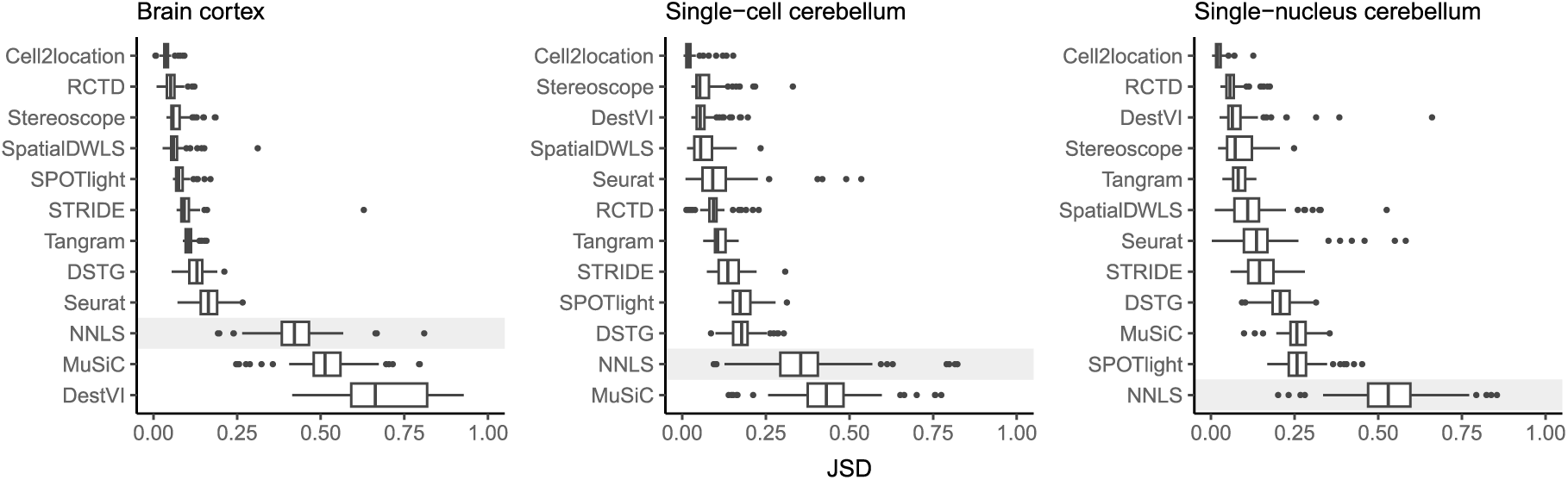
Prediction stability when using different reference datasets. For each synthetic dataset, we computed the Jensen-Shannon divergence between cell type proportions obtained from two different reference datasets.

### Evaluation of methods on public Visium datasets validate results on synthetic data

While synthetic datasets provide valuable benchmarks, it is crucial to validate deconvolution methods using real sequencing-based ST data, as they may exhibit distinct characteristics. To achieve this, we leveraged 10x Visium datasets from two mouse tissues, namely liver and skin cancer (melanoma). These datasets were chosen due to the availability of ground truth knowledge regarding cell proportions and localization, as elaborated further below.

### Liver

The liver is a particularly interesting case study due to its tissue pattern, where hepatocytes are highly abundant and constitute more than 60% of the tissue. This characteristic allows us to compare method performance with those of the *dominant cell type* abundance pattern from the silver standard, which was challenging for most methods. Here, we use the four Visium slides and single-cell dataset from the liver cell atlas of Guilliams et al. [34] (**Supplementary Table 1c**). The single-cell dataset was generated from three different experimental protocols– scRNA-seq following *ex vivo* digestion, scRNA-seq following *in vivo* liver perfusion, and single-nucleus RNA-seq (snRNA-seq) on frozen liver—additionally enabling us to assess method stability on different reference datasets.

We assessed method performance using AUPR and JSD by leveraging prior knowledge of the localization and composition of cell types in the liver. Although the true composition of each spot is not known, we can assume the presence of certain cell types in certain locations in the tissue due to the zonated nature of the liver. Specifically, we calculated the AUPR of portal vein and central vein endothelial cells (ECs) under the assumption that they are present only in their respective venous locations. Next, we calculated the JSD of the expected and predicted cell type proportions in a liver sample. The expected cell type proportions were based on snRNA-seq data (**Figure S11**), which has been shown to best recapitulate actual cell compositions observed by confocal microscopy [34]. We ran each method on a combined reference containing all three experimental protocols (*ex vivo* scRNA-seq, *in vivo* scRNA-seq, and snRNA-seq), as well as on each protocol separately. To ensure consistency, we filtered each reference dataset to only include the nine cell types that were common to all three protocols.

RCTD and cell2location were the top performers for both AUPR and JSD (**Figure 6a**), achieving a JSD comparable to those of biological variation, i.e., the average pairwise JSD between four snRNA-seq samples. In contrast, Tangram, DSTG, SPOTlight, and stereoscope had higher JSD values than those of NNLS. Except for SPOTlight, these three methods were not able to accurately infer the overabundance of the dominant cell type, with stereoscope and Tangram predicting a uniform distribution of cell types (**Figure S12**). This corresponds to what we observed in the *dominant cell type* pattern of the silver standard and certain FOVs in the gold standard. To further substantiate this, we used the *ex vivo* scRNA-seq dataset to generate 10 synthetic datasets with the *dominant cell type* abundance pattern. Then, we used the remaining two experimental protocols separately as the reference for deconvolution. Indeed, the method rankings followed the overall pattern, and a comparison of the JSD values between synthetic and real Visium data revealed a strong correlation (**Figure S13**).

**Figure 6.**
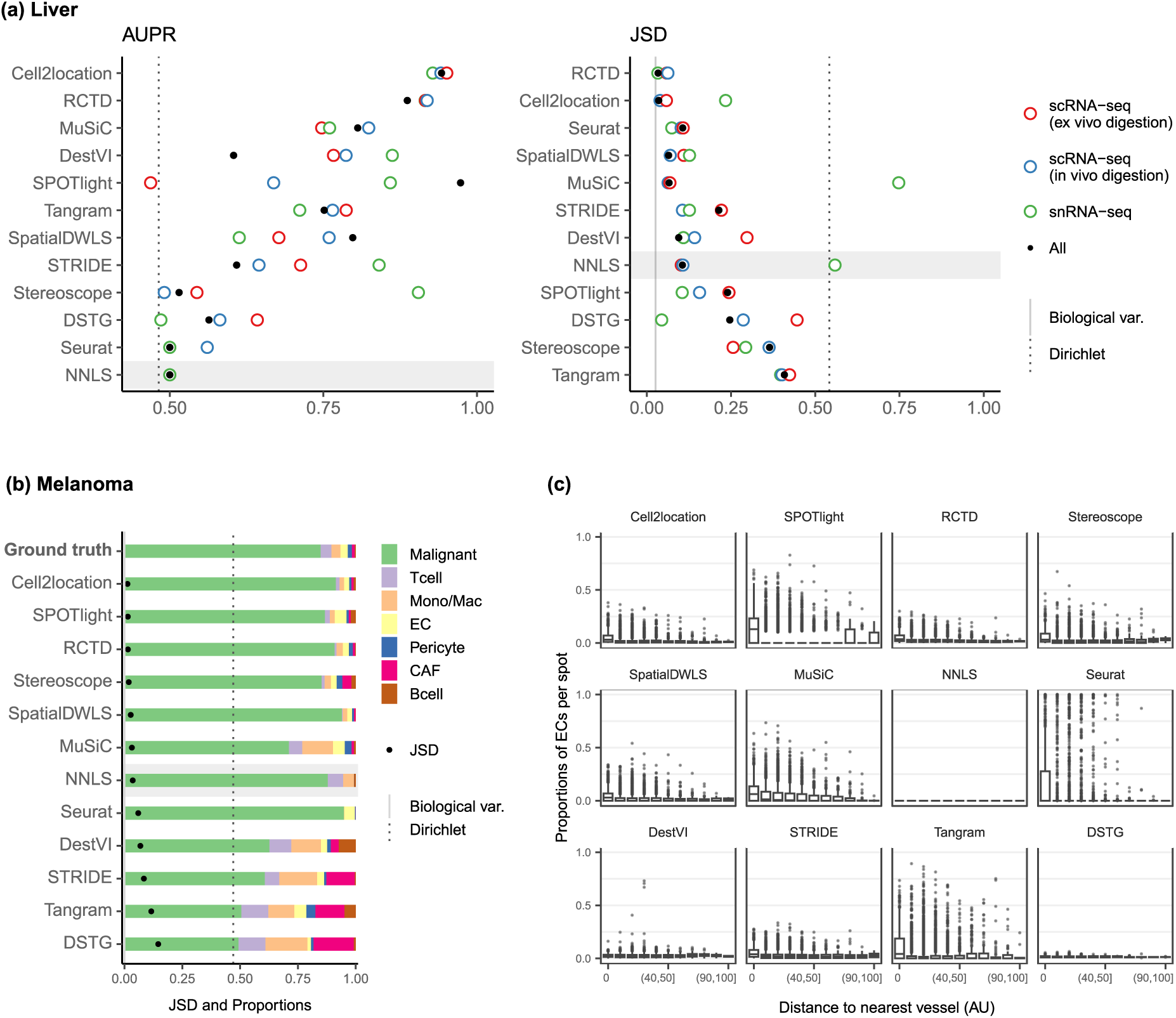
Method performance on two Visium case studies. **(a)** In the liver case study, the AUPR was calculated using the presence of portal/central vein endothelial cells in portal and central veins, and the JSD was calculated by comparing predicted cell type proportions with those from snRNA-seq. All reference datasets contain nine cell types. Biological variation refers to the average pairwise JSD between four snRNA-seq samples. Methods are ordered based on the summed rank of all data points. **(b)** For melanoma, the JSD was calculated between the predicted cell type proportions and those from Molecular Cartography (bold). Biological variation refers to the JSD between the two Molecular Cartography samples. **(c)** Relationship between the proportions of endothelial cells predicted per spot and their distance to the nearest blood vessel (in arbitrary units, AU), where zero denotes a spot annotated as a vessel. An inverse correlation can be observed more clearly in better-performing methods.

Nonetheless, there were some inconsistencies between the AUPR and JSD rankings, specifically with Seurat and SpatialDWLS ranking highly for JSD and lowly for AUPR. This is because the JSD is heavily influenced by the dominant cell type in the dataset, such that even when predicting only the dominant cell type per spot, Seurat still performed well in terms of JSD. However, it was unable to predict the presence of portal and central vein ECs in their respective regions (**Figure S14**). Therefore, complementary metrics like AUPR and JSD must both be considered when evaluating methods.

We observed that using the combination of the three experimental protocols as the reference dataset did not necessarily result in the best performance, and it was often possible to achieve better or similar results by using a reference dataset from a single experimental protocol. The best reference varied between methods, and most methods did not exhibit consistent performance across all references. Interestingly, Cell2location, MuSiC, and NNLS had much higher JSD when using the snRNA-seq data as the reference, while RCTD and Seurat had the lowest JSD on the same reference. To further evaluate the stability of the methods, we calculated the JSD between proportions predicted with different reference datasets. RCTD and Seurat showed the lowest JSD, indicating higher stability (**Figure S15**). Finally, we examined the predicted proportions when using the entire atlas without filtering cell types, which contains all three protocols and 23 cell types instead of only the nine common cell types (**Figure S12**). The additional 14 cell types made up around 20% of the ground truth proportions. While RCTD, Seurat, SpatialDWLS, and MuSiC retained the relative proportions of the nine common cell types, the rest predicted substantially different cell compositions.

### Melanoma

Melanoma poses a significant challenge in both therapeutic and research efforts due to its high degree of heterogeneity and plasticity. In a recent study, Karras et al. [35] investigated the cell state diversity of melanoma by generating single-cell and ST data from a mouse melanoma model (**Supplementary Table 1d**). Among others, the spatial data consists of three individual tumor sections profiled by 10x Visium, as well as 33 regions of interest from two tumor samples profiled by Molecular Cartography (MC), an imaging-based technology that can profile up to 100 genes at subcellular resolution. Using a custom processing pipeline, we obtained cell type annotations—and subsequently the cell type proportions— for each of the MC samples. These cell type proportions were consistent across different sections and samples, and were used as the ground truth for deconvolution (**Figure S16**). We aggregated the predicted proportions of the seven malignant cell states in the Visium slides, as we could not reliably annotate these cell states in the MC dataset.

To assess method performance, we calculated the average pairwise JSDs between the two MC samples and three Visium slides. Cell2location, SPOTlight, and RCTD were the best performers with JSDs of around 0.01 (**Figure 6b**). With the exception of SPOTlight performing well in this dataset, the rankings of the remaining methods followed the usual trend in the silver standards, with NNLS outperforming Seurat, DestVI, STRIDE, and DSTG. Additionally, we sought to corroborate these findings through a more qualitative approach by leveraging the blood vessel annotations provided in the original study. Given that by definition, endothelial cells form the linings of blood vessels, we visualized the relationship between EC abundance and the distance to the nearest vessel (**Figure 6c**). Although the majority of spots were predicted to have no ECs, a fraction of spots exhibited the expected trend where ECs were more abundant the closer the spot was to a blood vessel. This trend was more discernible for higher-ranked methods, while the lower-ranked ones either showed no correlation (NNLS, DestVI, DSTG) or had noisy predictions (Seurat, Tangram).

### Runtime and scalability

Most methods are able to deconvolve the silver standard datasets in less than 30 minutes and only had slight variability in the runtime (**Figure 7a**). Model-based methods—DestVI, stereoscope, cell2location, and STRIDE—have the advantage that a model built from a single-cell reference can be reused for all synthetic datasets derived from that reference (i.e., the nine abundance patterns × 10 replicates). This typically reduces the runtime needed to fit the model on the synthetic datasets. However, these methods tend to be more computationally intensive and are strongly recommended to be run with GPU acceleration. This was not implemented in STRIDE and could explain its longer runtime during model building.

**Figure 7.**
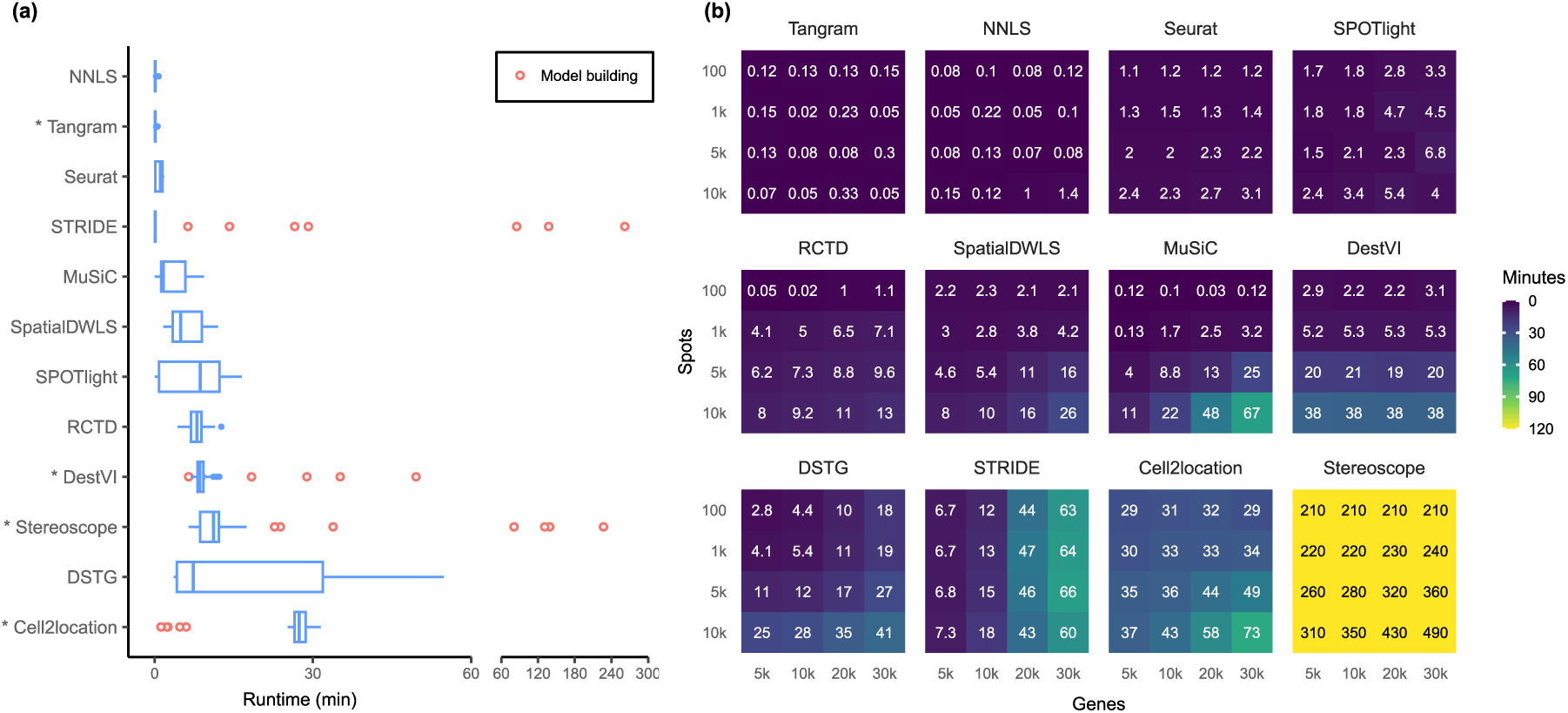
**(a)** Runtime over the 63 silver standards (three replicates each). Methods are ordered by total runtime. Asterisks indicate when GPU acceleration has been used. Cell2location, stereoscope, DestVI, and STRIDE first build a model for each single-cell reference (red points), which can be reused for all synthetic datasets derived from that reference. **(b)** Method scalability on increasing dimensions of the spatial dataset. For model-based methods, the model building and fitting time were summed. Methods are ordered based on total runtime.

Next, we investigated method scalability by varying the dimensions of the spatial dataset (**Figure 7b**). We tested 16 combinations of spots (100, 1k, 5k, 10k) and genes (5k, 10k, 20k, 30k), while the single-cell dataset was kept at 5k cells and 30k genes. Here, we considered both the model building and fitting step in the runtime (when applicable). Tangram, Seurat, and SPOTlight had a constant runtime across all dimensions, each deconvolving the largest dataset (3 x 10^8^ elements in total) in less than ten minutes. Other methods had increased runtime both when spots and genes were increased, except for DestVI which was only affected by the number of spots, and STRIDE by the number of genes. This was because by default, DestVI only uses 2,000 highly variable genes for deconvolution. Similarly, the scalability of RCTD, SpatialDWLS, Tangram, and SPOTlight are due to the fact that they only make use of differentially expressed or cell type-specific marker genes. Stereoscope was the least scalable by far, taking over eight hours for the largest dataset, and 3.5 hours for the smallest one. Note that many methods allow for parameters that can greatly affect runtime, such as the number of training epochs and/or the number of genes to use (**Supplementary Table 2**). For instance, here we have used all genes to run stereoscope (default parameters), but the authors have suggested that it is possible to use only 5,000 genes in the model and maintain the same performance.

## Discussion

In this study, we performed a thorough investigation of various biologically relevant and previously unconsidered aspects related to deconvolution. We evaluated the performance of 11 deconvolution methods on 63 silver standards, three gold standards, and two Visium datasets using three complementary metrics. We also incorporated two baselines for every analysis, the NNLS algorithm and null distribution proportions from a Dirichlet distribution. In the case studies, we demonstrated two approaches for evaluating deconvolution methods in datasets lacking an absolute ground truth. These approaches include using proportions derived from another sequencing or spatial technology as a proxy, and leveraging spot annotations, e.g., zonation or blood vessel annotations, that typically have already been generated for a separate analysis.

Our findings indicate that RCTD, cell2location, and SpatialDWLS were the highest ranked methods in terms of performance, consistent with previous studies [10], [11]. However, we also found that over half of the methods did not outperform the baseline (NNLS) and bulk deconvolution method (MuSiC). These results were consistent across the silver and gold standards, as well as the liver and melanoma case studies, demonstrating the generalizability and applicability of our simulation and benchmarking framework. We also found that the abundance pattern of the tissue and the reference dataset used had the most significant impact on method performance. Even top-performing methods struggled with challenging abundance patterns, such as when a rare cell type was present in only one region, or when a highly dominant cell type masks the signature of less abundant ones. Furthermore, different reference datasets could result in substantially different predicted proportions. Methods that accounted for technical variability in their models, such as cell2location and RCTD, were more stable to changes in the reference dataset than those that did not, such as SpatialDWLS.

Regarding the reference dataset, the number of genes per cell type (which is generally correlated to the sequencing depth) seems to have a significant impact on the performance of deconvolution methods. We observed that methods were less accurate in datasets with fewer genes per cell type (**Figure S9**). For example, all methods performed better in the snRNA-seq cerebellum dataset, which had the same number of cell types as the scRNA-seq cerebellum dataset, but on average 1,000 more genes per cell type. The kidney dataset was the most challenging for all methods, with most of its 16 cell types having less than 1,000 genes. This was evident from the RMSE and JSD scores that were relatively closer to the null distribution than in other datasets (**Figure S3,S5**). In contrast, the 18 cell types in the brain cortex dataset had an average of 3,000-9,000 features, leading to much better performance for most methods compared to the kidney dataset despite having more cell types. This trend was also observed in the STARMap gold standard dataset, which consisted of only 996 genes. Most methods performed worse in the STARMap dataset except for SpatialDWLS, SPOTlight, and Tangram. Since these three methods only use marker genes for deconvolution, this may explain why the small number of genes (most of which were already marker genes) did not affect them as much.

In addition to performance, the runtime and scalability of a method is also a factor to consider. Although most runtimes were comparable on our silver standards, we have made use of GPU acceleration for Tangram, DestVI, stereoscope, and cell2location. As this might not be available for some users, using these methods on the CPU might require training on fewer epochs or selecting only a subset of genes. With all these factors in consideration, we recommend RCTD as a good starting point. In addition to being one of the best and fastest methods, it also allows CPU parallelization (i.e., using multiple cores) for users that may not have access to a GPU.

As a general guideline, we recommend comparing the result of multiple deconvolution methods, especially between cell2location and RCTD. If the predictions are highly contradictory, the reference dataset may not be of sufficiently good quality. We recommend obtaining scRNA-seq and spatial data from the same sample to reduce biological variability, as there will always be technical variability across platforms. This can also ensure that the same cell types will be present in both datasets [9]. If that is not possible, use a reference dataset that has sufficient sequencing depth (at least more than 1,000 genes per cell type), preferably from a single platform. In addition to checking the sequencing depth, our simulator *synthspot* can also be used to evaluate the quality of the reference dataset. As we have demonstrated in the liver case study, users can generate synthetic spatial datasets with an abundance pattern that best resembles their tissue of interest. With a high-quality reference, both cell2location and RCTD should be able to achieve an AUPR close to one and JSD close to zero. Our Nextflow pipeline seamlessly integrates the complete workflow of synthetic data generation, deconvolution, and performance evaluation.

As spatial omics is still an emerging field, the development of new deconvolution methods can be anticipated in the future. Our benchmark provides a reliable and reproducible evaluation framework for current and upcoming deconvolution tools, making it a valuable resource for researchers in the field.

## Data availability

All datasets used in this article, including the silver standard, gold standard, and case studies are available on Zenodo at https://zenodo.org/records/10277187. Original download links and accession numbers of individual studies can be found at **Supplementary Table 1**.

## Code availability

The *synthspot* R package can be downloaded from https://github.com/saeyslab/synthspot. The Nextflow pipeline along with analysis scripts can be found at https://github.com/saeyslab/spotless-benchmark.

## Acknowledgements

We would like to thank all the authors of the methods who provided valuable feedback on how to optimally run their algorithms. Their input has been crucial in enabling us to compare the methods in a fair manner. We thank Lotte Pollaris for processing the Molecular Cartography dataset. This study was funded by BOF (Ghent University, C.S.), The Research Foundation – Flanders (1181318N and FWO SBO; iPSC LiMics, R.B.), the Flemish Government under the “Onderzoeksprogramma Artificiele Intelligentie (AI) Vlaanderen” programme (R.S.), BOF18-GOA-024 (Ghent University, Y.S.) and the Belgian Excellence of Science (EOS) program to Y.S.

## Methods

### Synthspot

We generated synthetic spatial datasets using the *synthspot* R package, whose spot generation process is modeled after that of SPOTlight [18]. This simulator generates synthetic spot data by considering the gene expression of one spot as a mixture of the expression of *n* cells, with *n* being a random number between 2 and 10 that is sampled from a uniform distribution. To generate one spot, the simulator samples *n* cells from the input scRNA-seq data, sums their counts gene by gene, and downsamples the counts. The target number of counts for downsampling is picked from a normal distribution with mean and standard deviation of 20,000 ± 5,000 by default, but these values can be changed by the user. To mimic biological tissue, *synthspot* generates artificial regions, or groups of spots with similar cell type compositions (**Figure S1**). The prior distribution of each cell type in each region is influenced by the selected *abundance pattern*, called *dataset type* in the package (**Supplementary Note 1**).

### Method execution

An overview of the 11 methods can be found in **Supplementary Table 2**. As input, all methods require a reference scRNA-seq dataset with cell type annotations along with a spatial dataset. We first ran the methods based on the provided tutorials, and later reached out to the original authors of each method for additional recommendations. For most methods, we explored different parameters and selected ones that resulted in the best performance. The specifics on how each method was run and the final parameters can be found in **Supplementary Note 3.** Unless stated otherwise, these parameters were applied on all datasets used in this study.

We implemented a Nextflow pipeline [36] to run the methods concurrently and compute their performance. For reproducibility, each method was installed inside a Docker container and can be run either using Docker or Apptainer (formerly known as Singularity). Our pipeline can be run in three modes: 1) *run_standard* to reproduce our benchmark, 2) *run_dataset* to run methods on a given scRNA-seq and spatial dataset, and 3) *generate_and_run* to also generate synthetic spatial data from the given scRNA-seq dataset. Our pipeline can accept both Seurat (.rds) and AnnData (.h5ad) objects as input, and the proportions are output as a tab-separated file. Parameter tuning has also been implemented.

All methods were deployed on a high-performance computing cluster with an Intel Xeon Gold 6140 processor operating at 2.3 GHz, running a RHEL8 operating system. We made use of GPU acceleration whenever possible using a NVIDIA Volta V100 GPU. We ran all methods with one core and 8 GB memory, dynamically increasing the memory by 8 GB if the process failed.

### Datasets

#### Gold

The seqFISH+ dataset consists of two mouse brain tissue slices (cortex and olfactory bulb) with seven field of views (FOVs) per slice [3]. Ten thousand genes were profiled for each FOV. Each FOV had a dimension of 206 μm x 206 μm and a resolution of 103 nm per pixel. We simulated Visium spots of 55-μm diameter and disregarded the spot-to-spot distance, resulting in nine spots per FOV and 126 spots for the entire dataset. The FOVs had varying number of cells and cell types per simulated spot (**Supplementary Table 1a**). We created a reference dataset for each tissue slice using the combined single-cell expression profiles from all seven FOVs. There were 17 cell types in the cortex tissue section and nine cell types in the olfactory bulb tissue section.

The STARMap dataset of mouse primary visual cortex (VISp) is 1.4 mm×0.3 mm and contains 1,020 genes and 973 cells [28]. We generated Visium-like spots as above, assuming that 1 pixel = 810 nm, as this information was not provided in the original paper. We removed any spot with hippocampal cell types, and removed unannotated and Reln cells from spots, resulting in 108 spots comprising 569 cells and 12 cell types in total (**Supplementary Table 1a**). We used the VISp scRNA-seq dataset from the Allen Brain Atlas as the reference [37]. After filtering for common genes, 996 genes remained.

#### Silver

We used six scRNA-seq datasets and one snRNA-seq dataset for the generation of silver standards (**Supplementary Table 1b; Figure S9**). All datasets are publicly available and already contain cell type annotations. We downsized each dataset by filtering out cells with an ambiguous cell type label and cell types with less than 25 cells, and kept only highly variable genes (HVGs) that are expressed in at least 10% of cells from one cell type. We set this threshold at 25% for the brain cortex dataset. We further downsampled the kidney dataset to 15,000 cells, and the melanoma dataset to 20,000 cells. For the two cerebellum datasets, we integrated them following the method of Stuart et al. [22], performed the same gene filtering approach on the integrated dataset, and only retained common cell types.

We generated the synthetic datasets using nine out of the 17 possible abundance patterns in *synthspot*. For each scRNA-seq dataset, we generated 10 replicates of each abundance pattern. For each replicate, we ran *generate_synthetic_visium* with five regions (*n_regions*), with minimally 100 spots and maximally 200 spots per region (*n_spots_min*, *n_spots_max)*, and target counts of 20,000 ± 5,000 per spot (*visium_mean,* and *visium_sd*). There were on average 750 spots per replicate.

#### Liver

We downloaded single-cell and spatial datasets of healthy mice from the liver cell atlas [34] (**Supplementary Table 1c**). The single-cell data contains 185,894 cells and 31,053 genes from three experimental protocols: scRNA-seq following *ex vivo* digestion, scRNA-seq following *in vivo* liver perfusion, and snRNA-seq on frozen liver. We used the finer annotation of CD45^-^ cells, which among others, subclustered endothelial cells into portal vein, central vein, lymphatic, and liver sinusoidal endothelial cells. We only retained cell types where at least 50 cells are present in each protocol, resulting in nine common cell types. The spatial data consisted of four Visium slides, containing on average 1,440 spots. Each spot has been annotated by the original authors as belonging to either the central, mid, periportal, or portal zone. This zonation trajectory was calculated based on hepatocyte zonation markers.

We deconvolved the Visium slides using five variations of the single-cell reference: the entire dataset, the dataset filtered to nine cell types, and each experimental protocol separately (also filtered to nine cell types). The proportions obtained from using the entire dataset was not used to compute evaluation metrics but only to visualize method stability (**Figure S12**). Due to the large size of the reference, we ran some methods differently. We ran stereoscope with 5,000 HVGs and subsampled each cell type to a maximum of 250 cells (*-sub*). We gave STRIDE the number of topics to test (*--ntopics 23 33 43 53 63*) instead of the default range that goes up to triple the number of cell types. We ran DestVI with 2500 training epochs and batch size of 128. For Seurat, we ran *FindTransferAnchors* with the “rpca” (reciprocal PCA) option instead of “pcaproject”. MuSiC and SpatialDWLS use dense matrices in their code (in contrast to sparse matrices in other methods), resulting in a size limit of 2^31^ elements in a matrix. Whenever this is exceeded, we downsampled the reference dataset by keeping maximally 10,000 cells per cell type and keeping 3,000 HVGs along with any genes that are at least 10% expressed in a cell type. When a reference with multiple protocols was used, we provided this information to cell2location and Seurat.

To compute the AUPR, we considered only spots annotated as being in the central or portal zone. We considered the following ground truth for portal vein and central vein ECs:

|  | Portal vein EC | Central Vein EC |
| --- | --- | --- |
| Portal spot | 1 | 0 |
| Central spot | 0 | 1 |

For the JSD ground truth, we only retained samples from the snRNA-seq protocol where all nine cell types were present, resulting in four samples (ABU11, ABU13, ABU17, and ABU20). For both the ground truth and predicted proportions, we averaged the abundance of each cell type in per slide or sample. Then, we calculated the pairwise JSD between each of the four slides and four snRNA-seq samples and reported the average JSD. The biological variation was obtained by averaging the pairwise JSD between the snRNA-seq samples.

#### Melanoma

The scRNA-seq and spatial datasets were downloaded from the original study [35] (**Supplementary Table 1d**), with the scRNA-seq dataset being the same one used in the silver standard before the split (**Supplementary Table 1b**). We preprocessed the Visium slides by filtering out spots with fewer than 100 features, as their inclusion led to errors when running marker-based deconvolution methods. This filtering removed 26 spots from the third slide. The blood vessel annotation was provided by the pathologist from the original publication. We ran stereoscope and DestVI with the same parameters as the liver case study due to the size of the reference dataset.

The Molecular Cartography (MC) dataset consists of two samples, six sections, and 33 regions of interest (ROI). For each ROI, a DAPI-stained image and a text file containing transcript locations are provided. We obtained cell type proportions using an in-house image processing pipeline. First, the image was cleaned by eliminating tiling effects and background debris using BaSic [38] and scikit-image [39], respectively. Subsequently, DAPI-stained images were segmented, and locations of individual nuclei were obtained using CellPose [40]. The transcripts of all measured genes were then assigned to their corresponding cells, resulting in the creation of cell-by-gene count matrices. These matrices were normalized based on the size of segmented nuclei and preprocessed in ScanPy [41]. Specifically, counts were log-transformed, scaled, and genes expressed in fewer than five cells and cells with less than 10 transcripts were filtered out. Leiden clustering [42] was performed using 17 principal components and 35 neighbors, and cells were annotated using *scanpy.tl.score_genes* with a curated marker gene list. Finally, the counts of each cell type were aggregated across samples to obtain cell type proportions.

As dendritic cells and melanoma cell states could not be annotated in the MC dataset, we adjusted the predicted proportions from Visium by removing dendritic cells proportions (pDCs and DCs) and recalculating the relative proportions, and aggregating the proportions of the seven malignant cell states.

### Scalability

We generated a synthetic dataset with 10,000 spots and 31,053 genes using only the snRNA-seq protocol from the liver atlas. We then used the remaining two protocols (combined) as the reference dataset. Genes of the synthetic dataset were downsampled randomly based on a uniform distribution, while genes of the reference data were downsampled based on marker genes and HVGs. The spots/cells of both datasets were downsampled randomly, and this was done in a stratified manner for the reference dataset.

### Evaluation metrics and baselines

The root-mean-squared error between the known and predicted proportions (*p*) of a spot *s* for cell type *z*, in a reference dataset with *Z* cell types in total, is calculated as

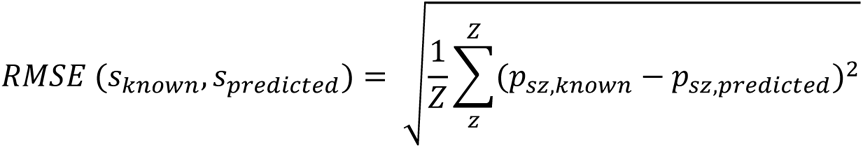

We calculated the JSD and AUPR using the R packages *philentropy* and *precrec*, respectively [43], [44]. The JSD is a smoothed and symmetric version of the Kullback-Leibler divergence (KL). It is calculated as

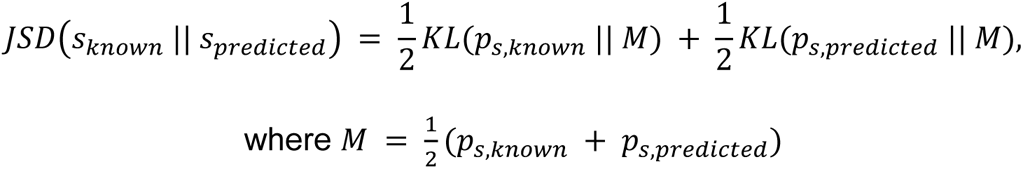

To calculate the AUPR, we binarized the known proportions by considering a cell type to be present in a spot if its proportion is greater than zero, and absent if it is equal to zero. Then, we compute the micro-averaged AUPR by aggregating all cell types together.

For the silver standards, the RMSE and JSD values across all spots are averaged and used as the representative value for a replicate *k* for each dataset-abundance pattern combination. In contrast, only one AUPR value is obtained per replicate.

We created the performance summary plot using the *funkyheatmap* package in R. To aggregate per data source and per abundance pattern, we calculated the grand mean of each metric across datasets, applied min-max scaling on each metric, and then computed the geometric mean of the scaled metrics (RMSE, AUPR, and JSD for gold and silver standards; AUPR and JSD for liver; and only JSD for melanoma). The aggregation per metric is the arithmetic mean of all datasets evaluated on that metric. Finally, we determined the overall rankings based on the weighted ranks of the following criteria: silver standard, gold standard, liver case study, melanoma case study, rare cell type detection, stability, and runtime. We assigned a weight of 0.5 to each silver standard abundance pattern and a weight of one to the remaining criteria.

To provide reference values for each metric, we used 1) random proportions based on probabilities drawn from a Dirichlet distribution and 2) predictions from the NNLS algorithm. For the former, we used the *DirichletReg* package [45] to generate reference values for all 63 silver standards using the average value across 100 iterations. The dimension of the α vector was equal to the number of cell types in the corresponding dataset, and all concentration values were set to one. For the NNLS baseline, we used the Lawson-Hanson NNLS implementation from the *nnls* R package. We solve for β in the equation **Y** = **X**β, with **Y** the spatial expression matrix and **X** the average gene expression profile per cell type from the scRNA-seq reference. We obtained proportion estimates by dividing each element of β with its total sum.

