## Supplementary Figures for "Spotless: a reproducible pipeline for benchmarking cell type deconvolution in spatial transcriptomics"

**(a)**

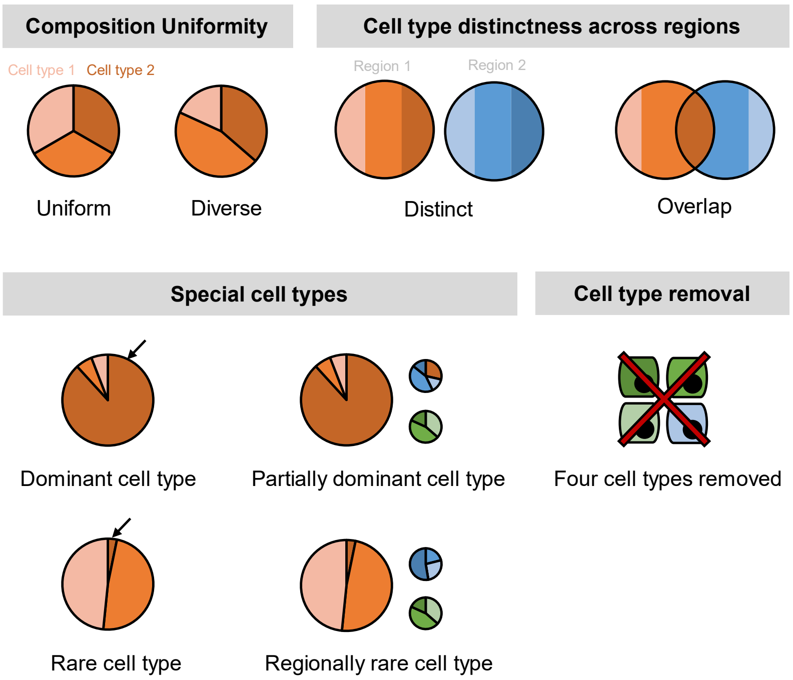


**(b)**

| Abundance patterns | Composition uniformity | Region Distinctness | Special cell types | Cell type removal? |
|---|---|---|---|---|
| **uniform distinct** | Uniform | Distinct | | |
| **uniform overlap** | Uniform | Overlap | | |
| **diverse distinct** | Diverse | Distinct | | |
| **diverse overlap** | Diverse | Overlap | | |
| **dominant** | Diverse | Overlap | Dominant | |
| **partially dominant** | Diverse | Overlap | Partially dominant | |
| **rare** | Diverse | Overlap | Rare | |
| **regional rare** | Diverse | Overlap | Regionally rare | |
| **missing cell type** | Diverse | Overlap | | Yes |

**Figure S1. (a)** Characteristics considered in synthspot include composition uniformity, cell type distinctness across regions, the presence of special cell types, or the removal of certain cell types. The pie charts represent frequency priors per region. Cell type composition in each region can be either *uniform* or *diverse,* depending on whether each cell type has the same or different number of cells per spot. *Distinct* and *overlap* refers to whether each region has a distinct set of cell types per region, or whether cell types can be found in multiple regions. *Dominant cell type* means that there is one dominant cell type that is 5-15 times more abundant than other cell types in all regions. In *partially dominant cell type*, this dominant cell type is dominant in all regions except for a region where it is equally abundant with other cell types and another region where it is absent. *Rare cell type* is the opposite of the dominant cell type, where one cell type is instead much less abundant than other cell types in all regions. In the *regional rare cell type* dataset, the rare cell type is only present in one region instead of all regions. **(b)** Each abundance pattern is a combination of multiple characteristics.

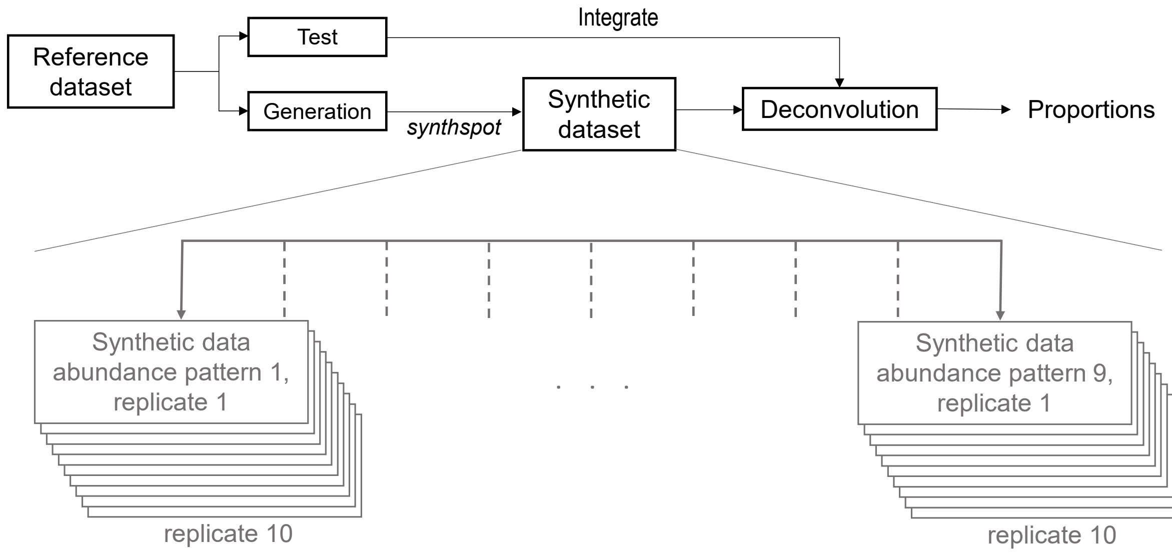


**Figure S2.** Silver standards are generated using half of the cells from a reference scRNA-seq dataset (generation). The other half (test) is used as the reference profile in deconvolution methods. This split is stratified by cell type. One scRNA-seq dataset gives rise to 90 synthetic datasets, as we generate nine *synthspot* abundance patterns with 10 replicates for each abundance pattern.

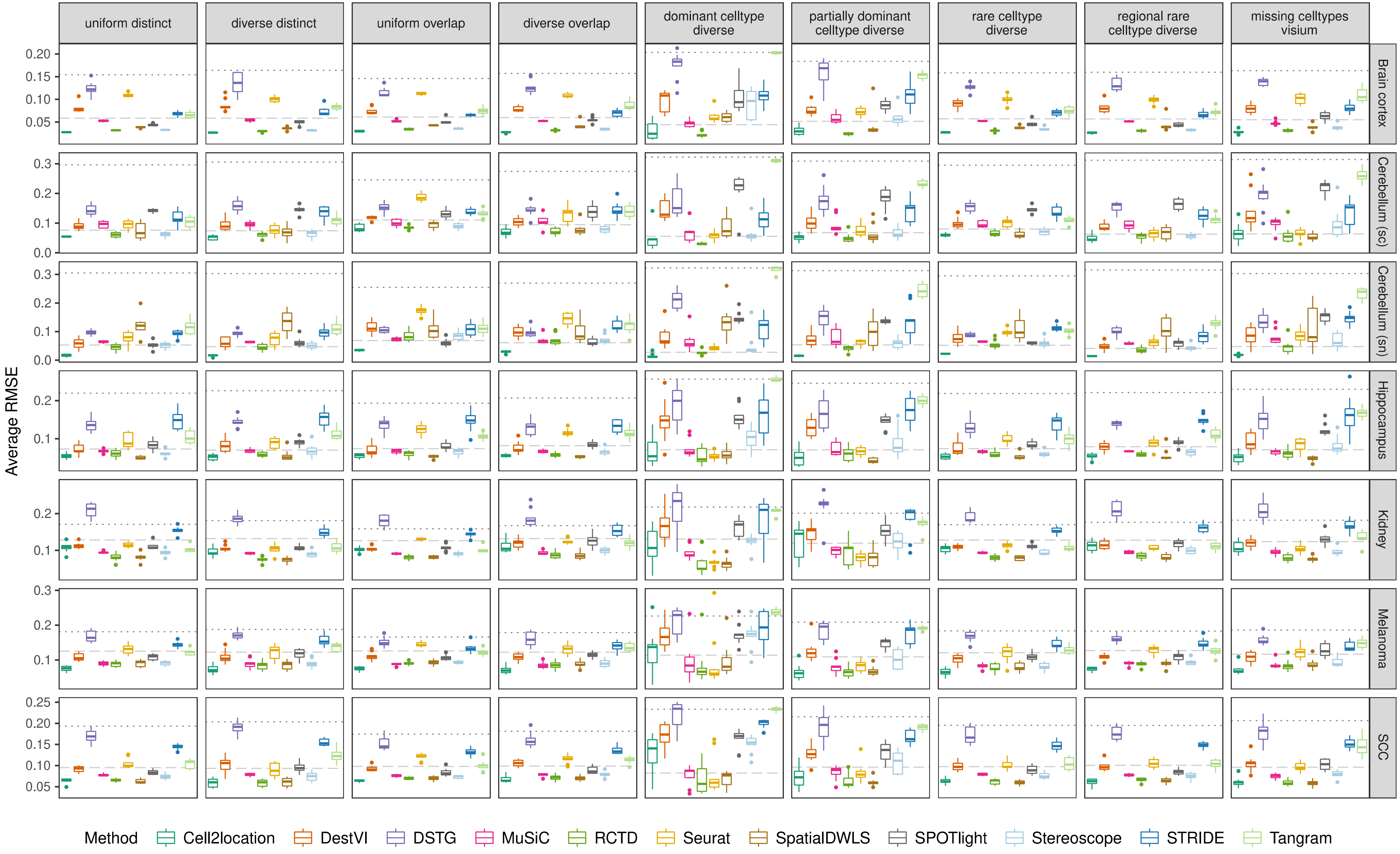


**Figure S3.** Boxplots of root-mean-squared error (RMSE) across ten replicates for each silver standard dataset (row) and abundance pattern (column). Long dashed gray line: baseline algorithm (non-negative least squares); dotted gray line: null distribution (random proportions from a Dirichlet distribution). Number of cell types: brain cortex, 18; single-cell cerebellum, 8; single-nucleus cerebellum, 8; hippocampus, 12; kidney, 16; melanoma, 15; SCC, 14.

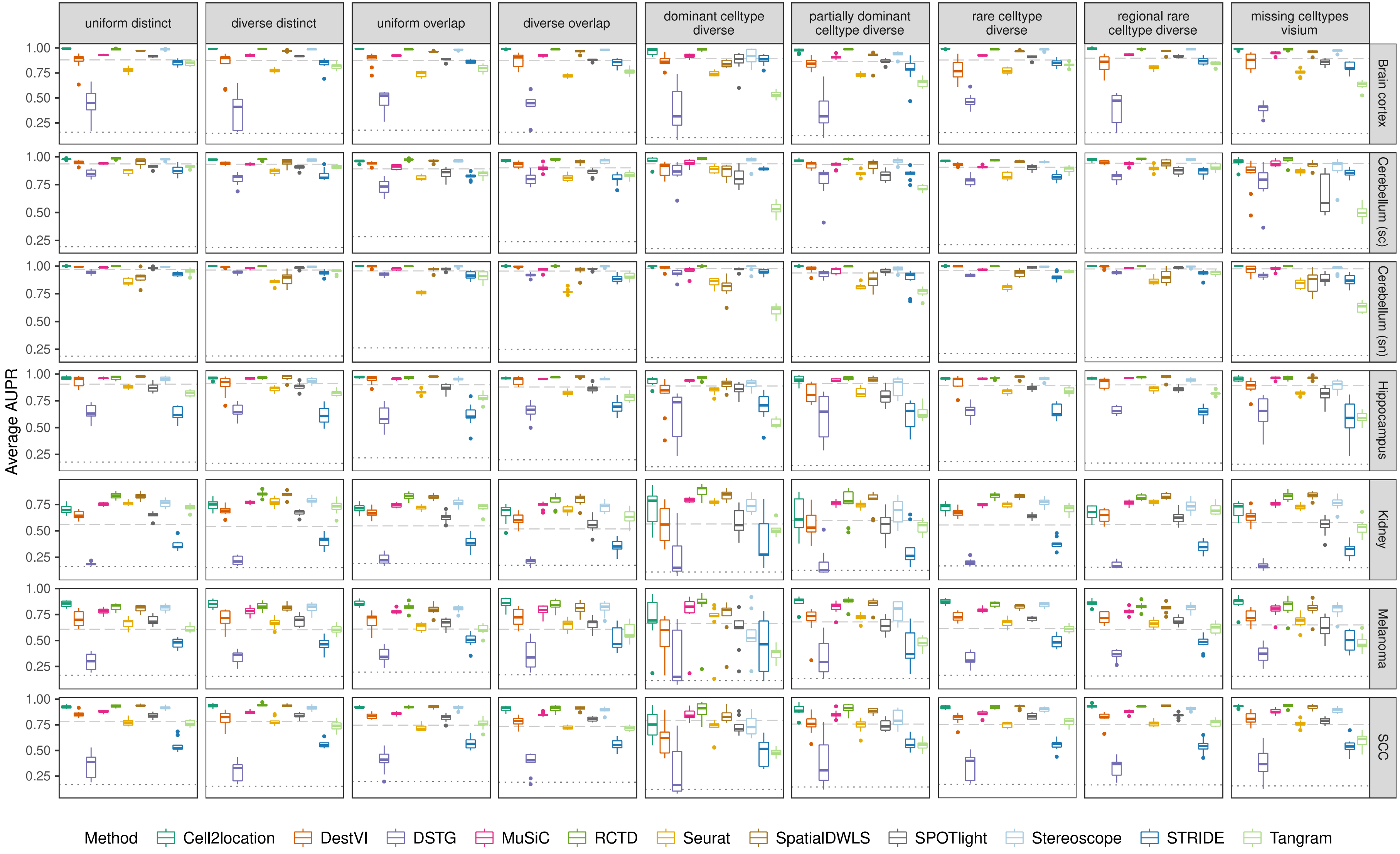


**Figure S4.** Boxplots of the area under the precision-recall curve (AUPR) across ten replicates for each silver standard dataset (row) and abundance pattern (column). Long dashed gray line: baseline algorithm (non-negative least squares); dotted gray line: null distribution (random proportions from a Dirichlet distribution). Number of cell types: brain cortex, 18; single-cell cerebellum, 8; single-nucleus cerebellum, 8; hippocampus, 12; kidney, 16; melanoma, 15; SCC, 14.

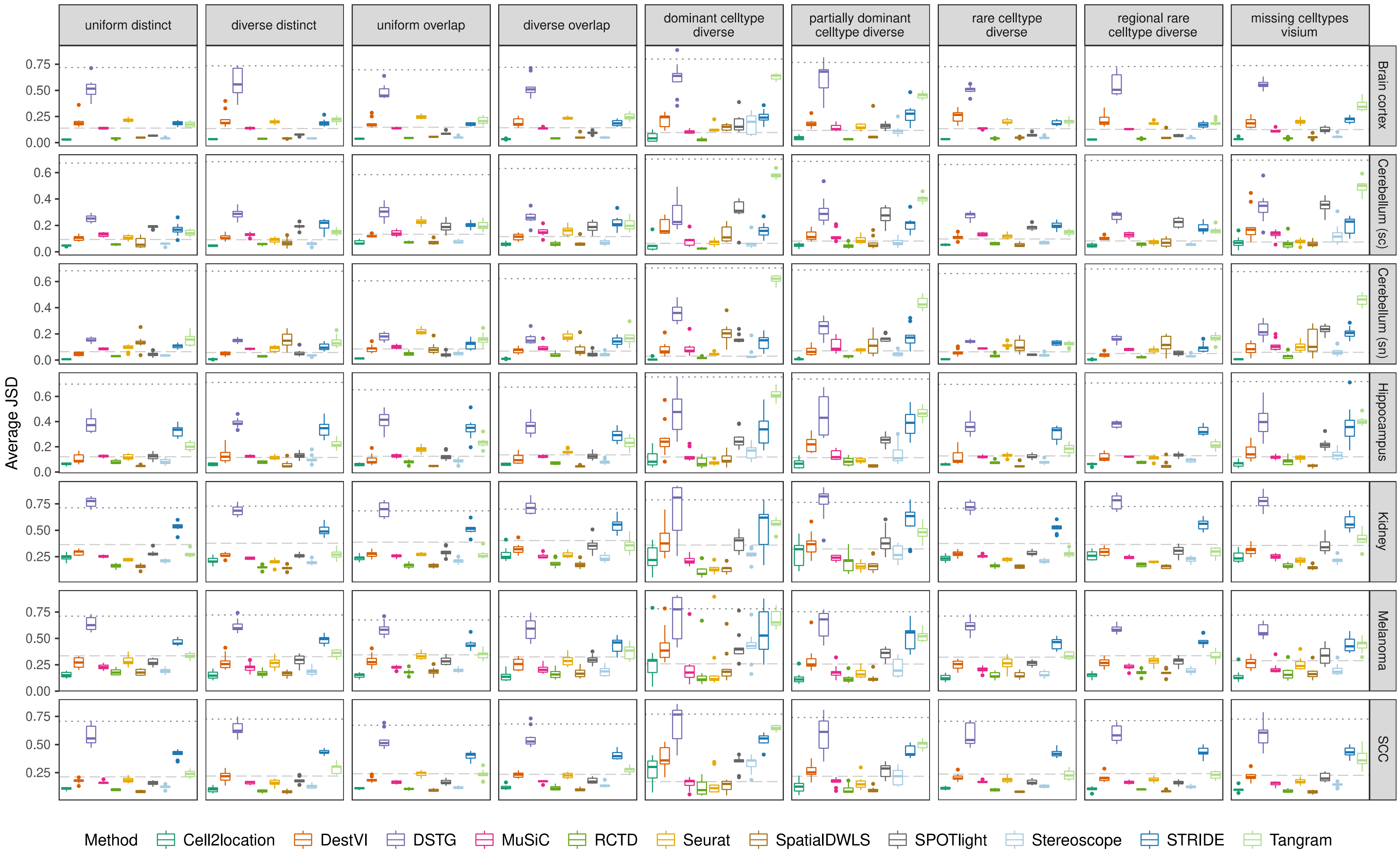


**Figure S5.** Boxplots of Jensen-Shannon divergence across ten replicates for each silver standard dataset (row) and abundance pattern (column). Long dashed gray line: baseline algorithm (non-negative least squares); dotted gray line: null distribution (random proportions from a Dirichlet distribution). Number of cell types: brain cortex, 18; single-cell cerebellum, 8; single-nucleus cerebellum, 8; hippocampus, 12; kidney, 16; melanoma, 15; SCC, 14.

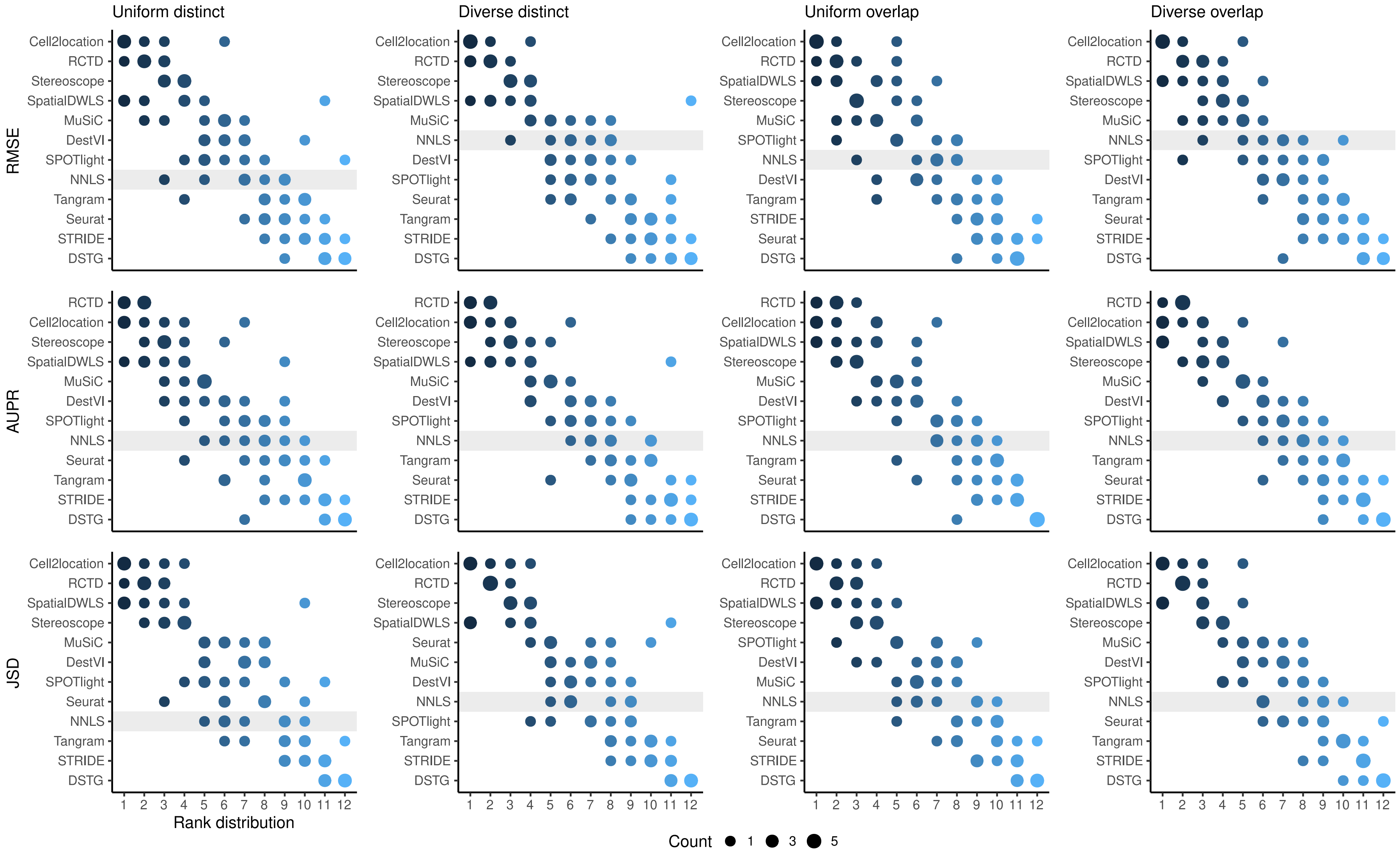


**Figure S6.** Summed rank plots across all silver standard datasets for each abundance pattern (column) and metric (row). Methods are ordered from best to worst performance. The baseline method (non-negative least squares) is shaded in gray.

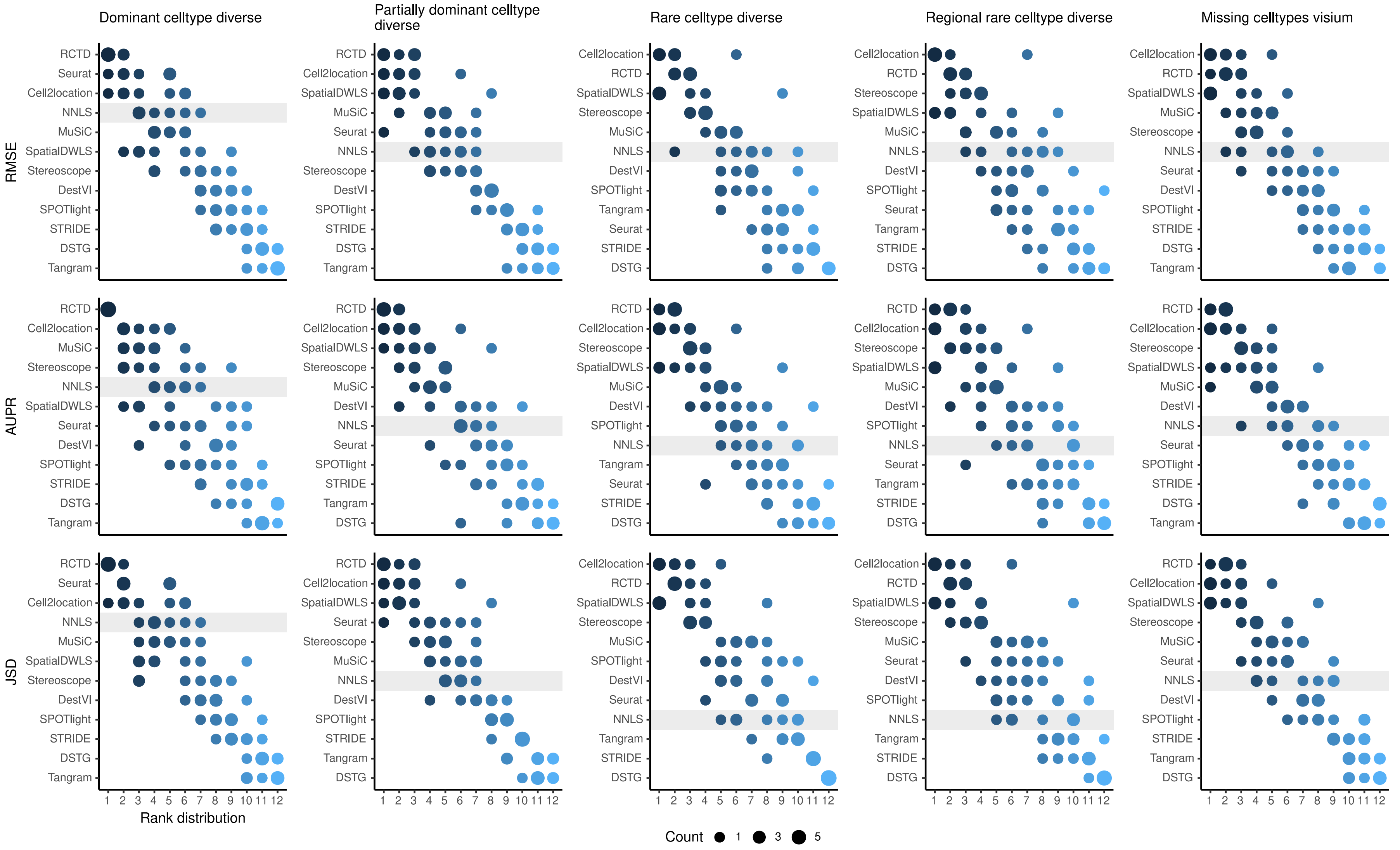


**Figure S6 (cont).** Summed rank plots across all silver standard datasets for each abundance pattern (column) and metric (row). Methods are ordered from best to worst performance. The baseline method (non-negative least squares) is shaded in gray.

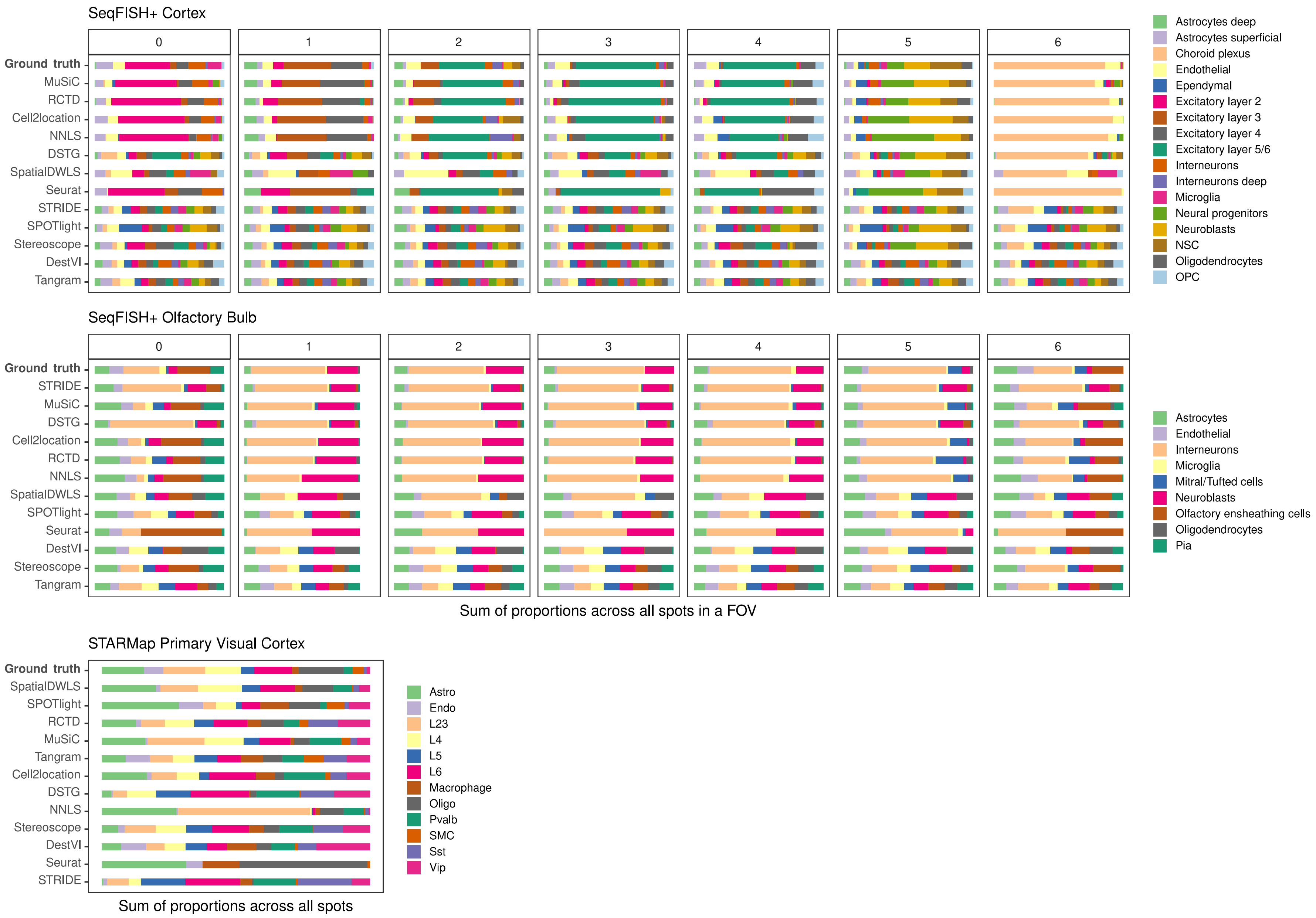


**Figure S7.** Summed abundances across all spots for each gold standard dataset. FOV1 of the olfactory bulb dataset has shorter bars as it only contains 8 spots. Methods are ordered according to their ranking for each dataset, based on the lowest median RMSE.

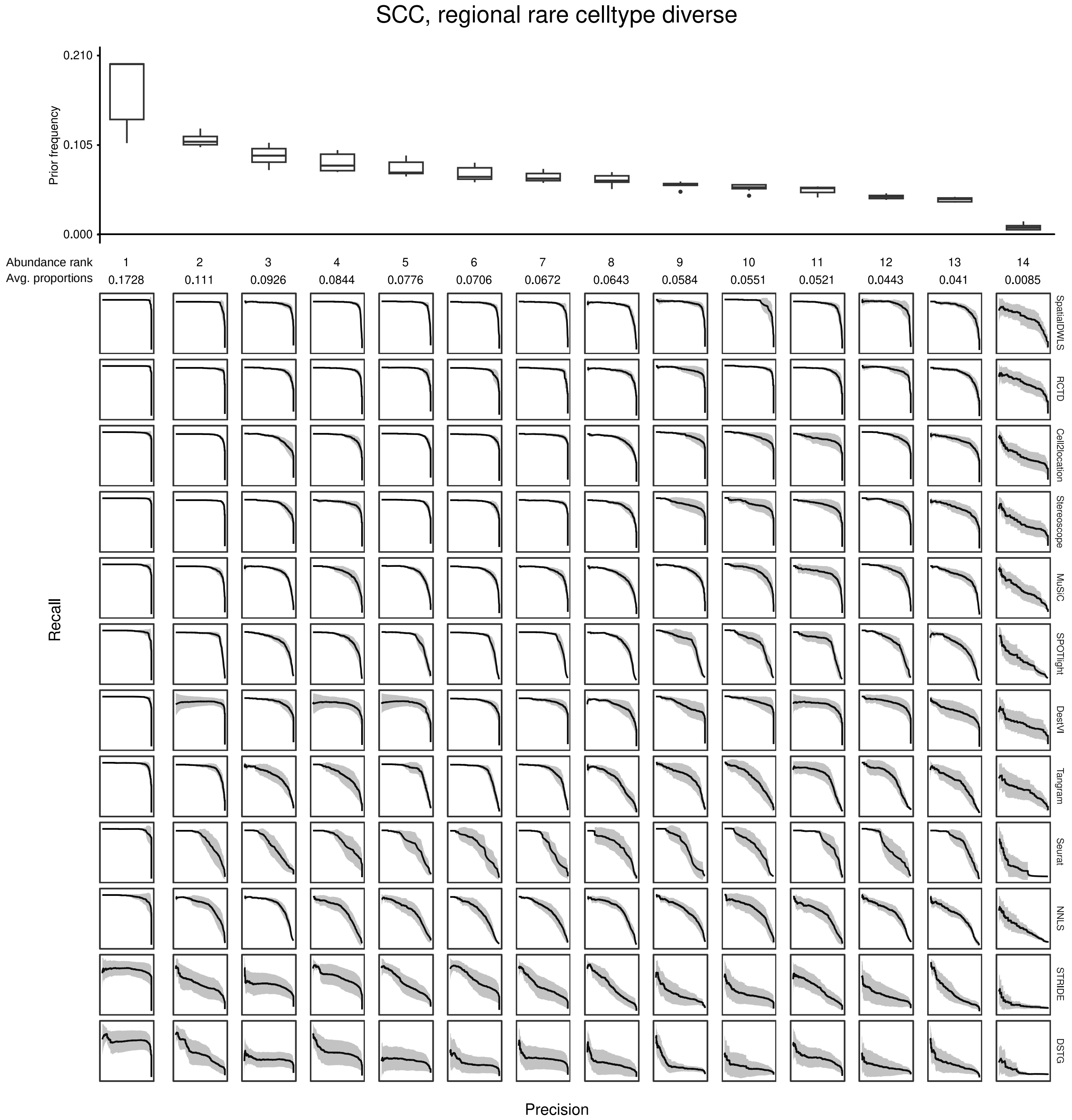


**Figure S8.** Evaluating the AUPR as a function of cell type abundance. Across the ten replicates of a silver standard dataset (here, the SCC dataset with the *regionally rare* abundance pattern is shown), we ordered cell types according to their abundance and computed the average precision-recall curve for each abundance rank. For instance, the leftmost column depicts the average precision-recall curve for the most abundant cell type in all ten replicates. The boxplots (top) depict cell type frequency priors in each abundance rank, i.e., the likelihood in which a cell type will be sampled in a spot. The observed average abundance is also indicated underneath the abundance rank number. As cells become less abundant, they become harder to detect by all methods, as indicated by less confident PR curves with smaller areas. Plots of all silver standard datasets can be found in our GitHub repository.

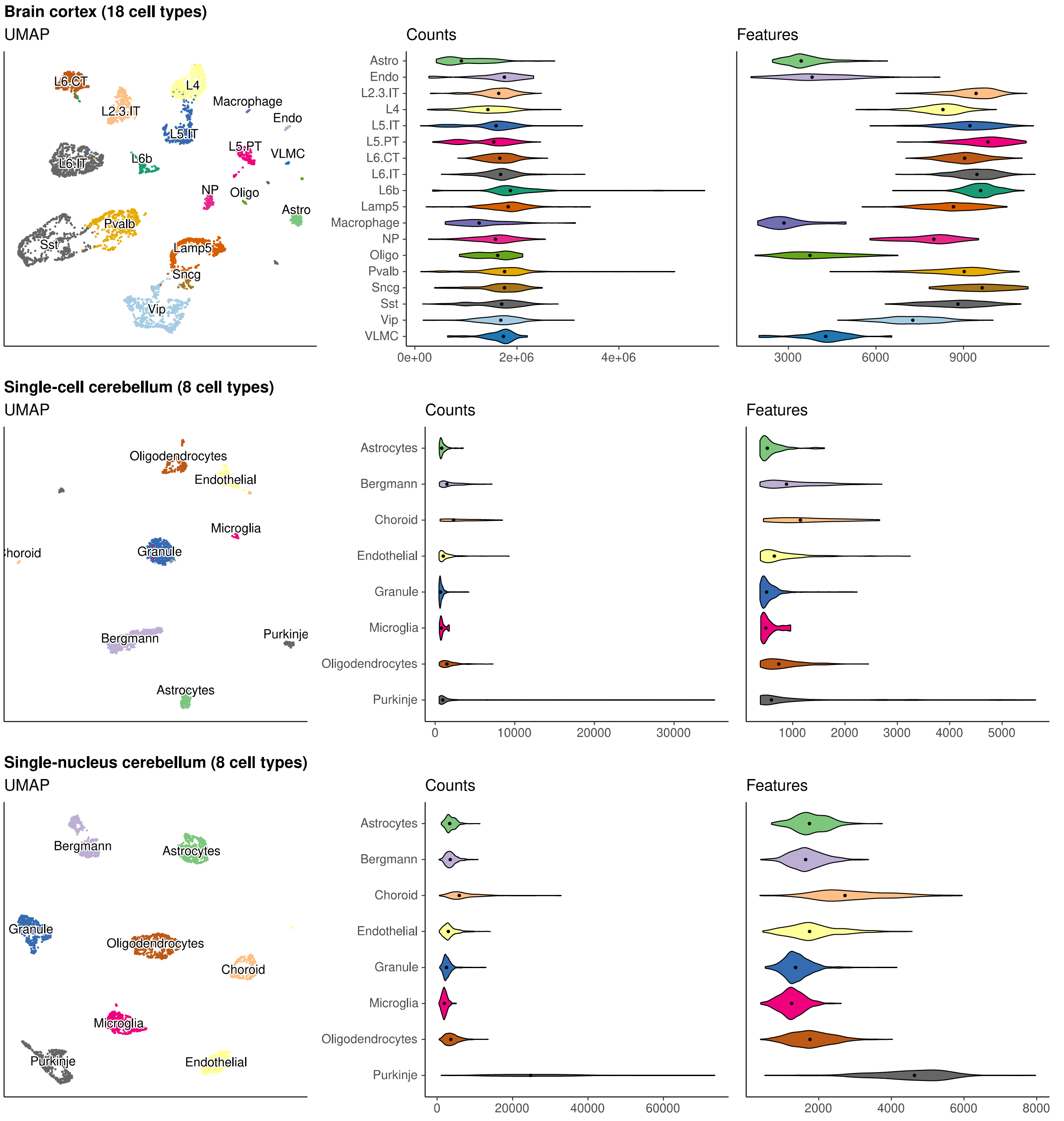


**Figure S9.** UMAP and violin plots of the seven scRNA-seq datasets used to generate silver standards. Each point on the UMAP represents a single cell and the distance between two points corresponds to how similar their gene expression profiles are. The violin plots show the distribution of total number of counts and features (genes) across all cells in a cell type. The brain cortex dataset has much higher counts than the others because it was sequenced with a plate-based method (SMART-Seq), while the others were sequenced with droplet-based methods (10x Chromium and Drop-seq).

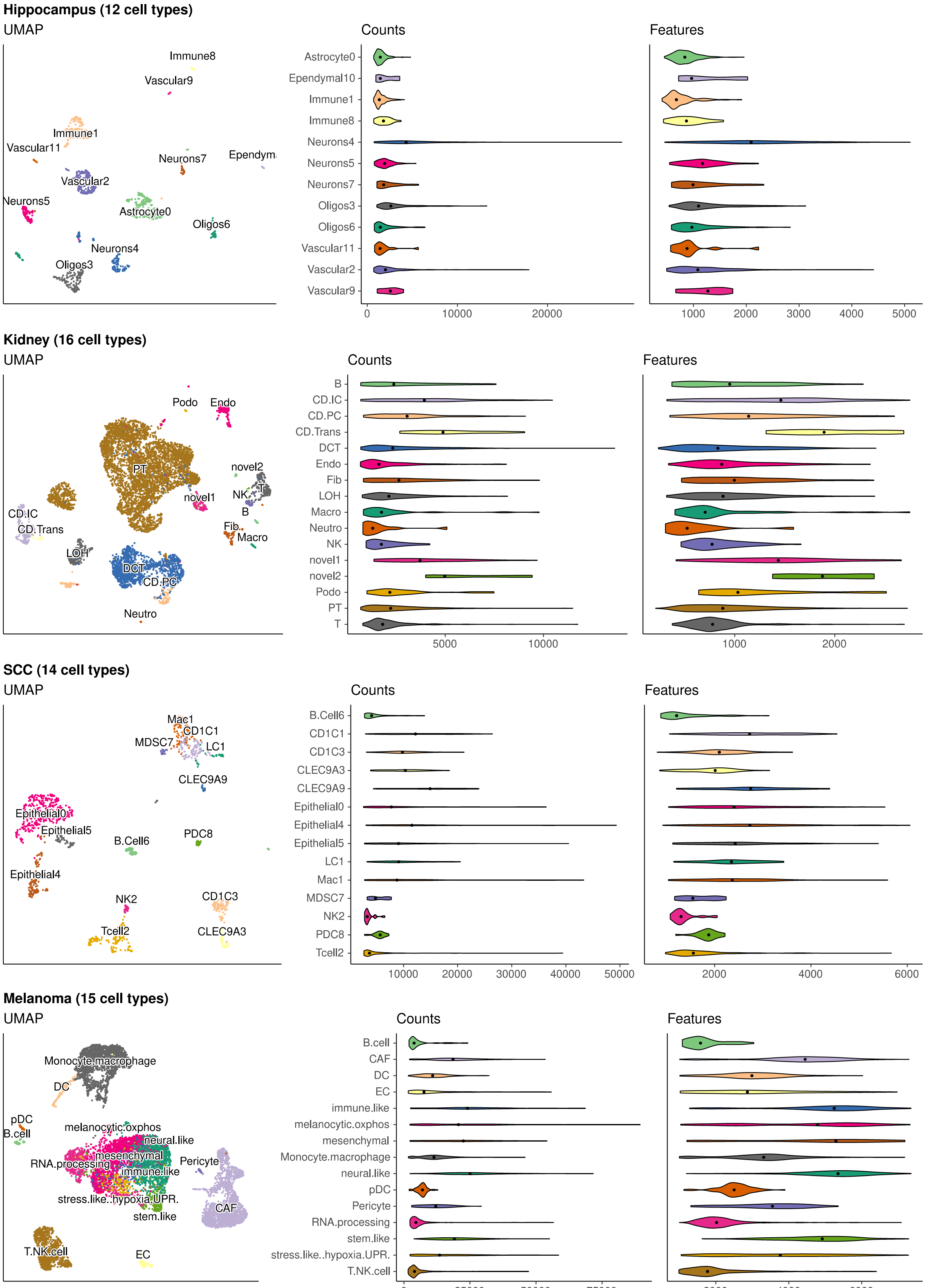


**Figure S9 (cont).** UMAP and violin plots of the seven scRNA-seq datasets used to generate silver standards. Each point on the UMAP represents a single cell and the distance between two points corresponds to how similar their gene expression profiles are. The violin plots show the distribution of total number of counts and features (genes) across all cells in a cell type. The brain cortex dataset has much higher counts than the others because it was sequenced with a plate-based method (SMART-Seq), while the others were sequenced with droplet-based methods (10x Chromium and Drop-seq).

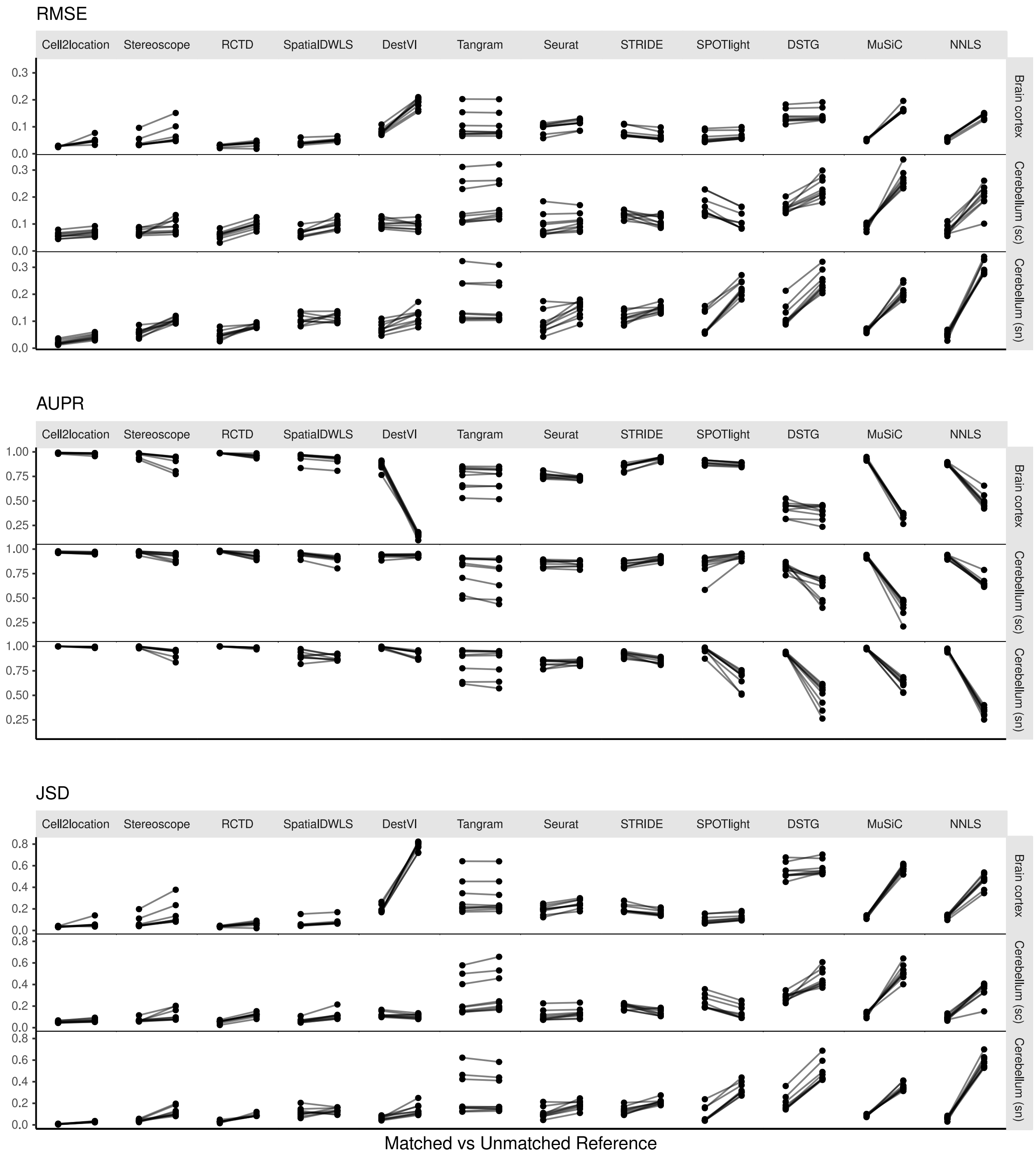


**Figure S10.** Changes in performance metrics when using a different reference dataset, from a matched to an unmatched reference (i.e., intra-dataset vs inter-dataset scenario). A matched reference means that the simulated spots and reference were generated from the same scRNA-seq data but on different halves of cells (**Figure S2**). Methods are ordered based on JSD between predicted proportions and not on the metrics themselves (**Figure 5**).

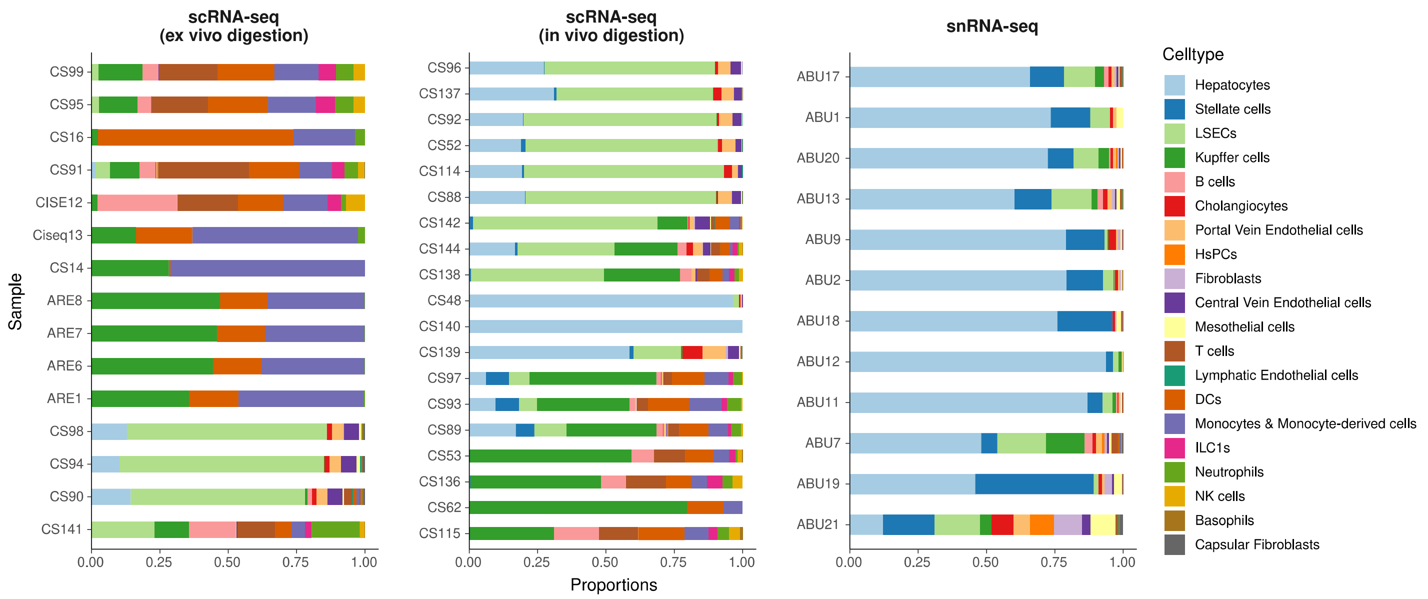


**Figure S11.** Comparison of cell type compositions between three sequencing protocols in the mouse liver atlas from Guilliams et al. (2022). In the original paper, samples profiled by snRNA-seq were shown to best resemble *in vivo* cell compositions.

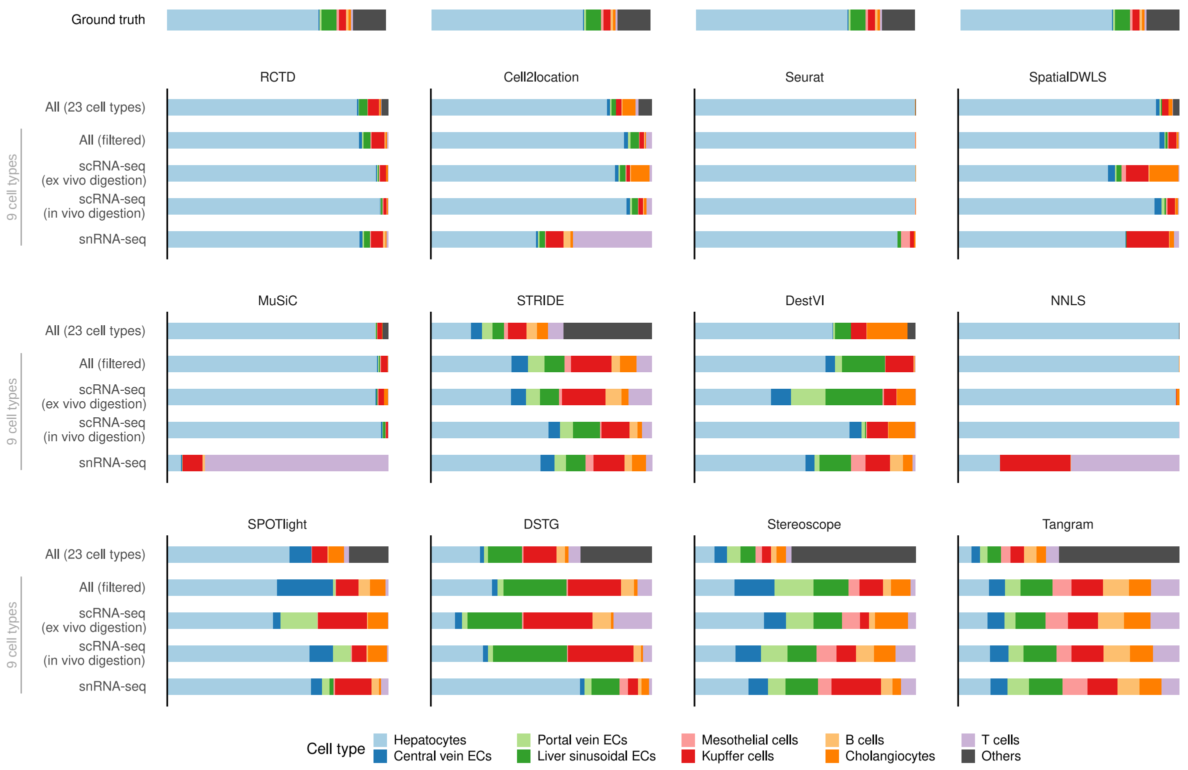


**Figure S12.** Predicted cell type abundances averaged across all four Visium slides from the liver atlas. Using different reference datasets for deconvolution often led to drastically different predictions (most notably with snRNA-seq). Each reference dataset except "All" was filtered to only contain nine common cell types. "All (filtered)" comprises all three protocols but only nine cell types; "All" contains 14 more cell types which were grouped under "Others". The ground truth is the average of four snRNA-seq samples (ABU11, ABU13, ABU17 ABU20). Methods are ordered according to their JSD rankings (**Figure 6a**).

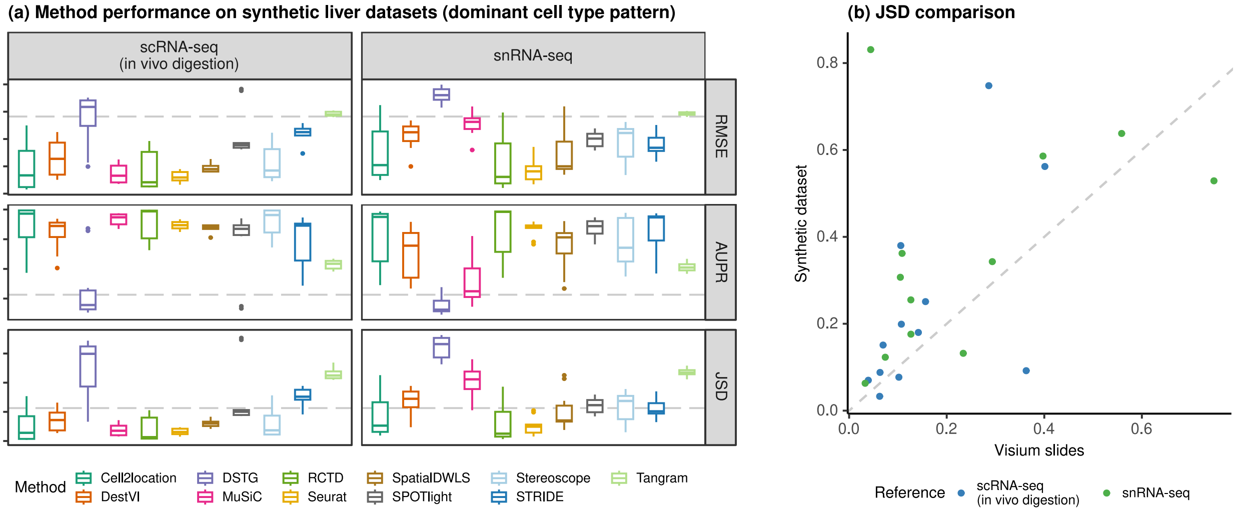


**Figure S13. (a)** Method performance on synthetic liver datasets generated from the *ex vivo* scRNA-seq protocol, using either the *in vivo* scRNA-seq or snRNA-seq protocols as the reference for deconvolution. **(b)** Comparison of the average JSD of each method between the synthetic and Visium datasets.

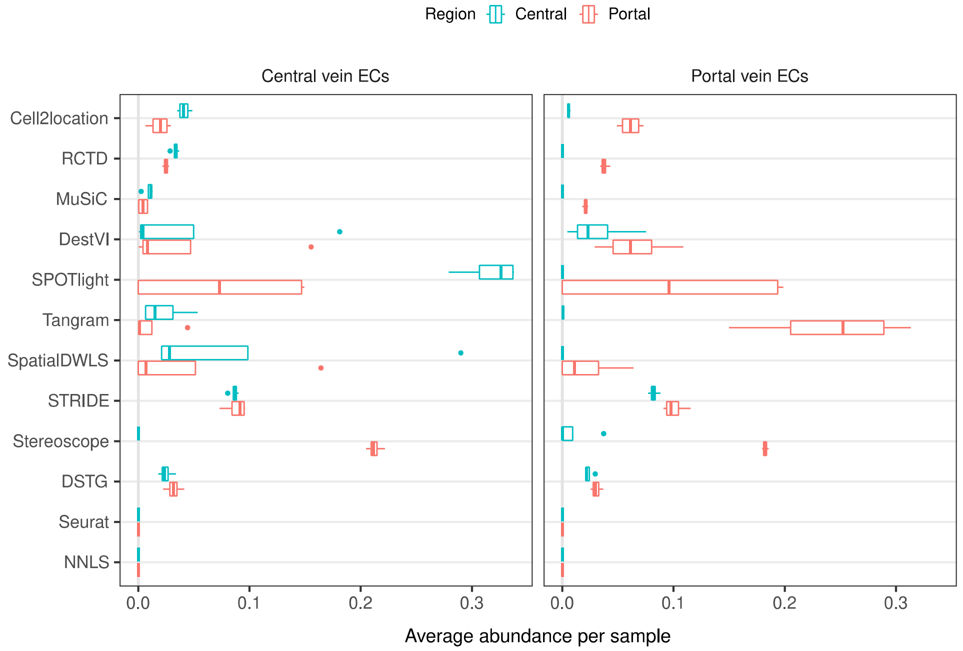


**Figure S14.** The predicted abundance of central vein and portal vein endothelial cells (ECs) for each spot in one liver Visium slide, using the reference dataset with all protocols but filtered to nine cell types. According to prior knowledge, central vein ECs should only be present in the central vein, and portal vein ECs in the portal vein. Methods are ordered based on their overall AUPR rankings (**Figure 6a**).

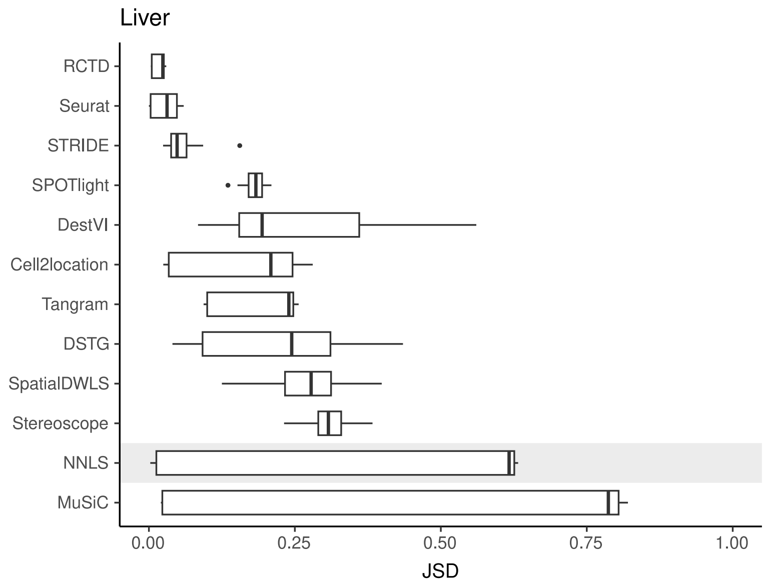


**Figure S15.** Stability of predicted proportions when using three different protocols from the liver atlas as reference for deconvolution. For each Visium slide, pairwise JSD values between each reference were calculated. Methods are ordered based on stability, with a lower JSD indicating higher stability.

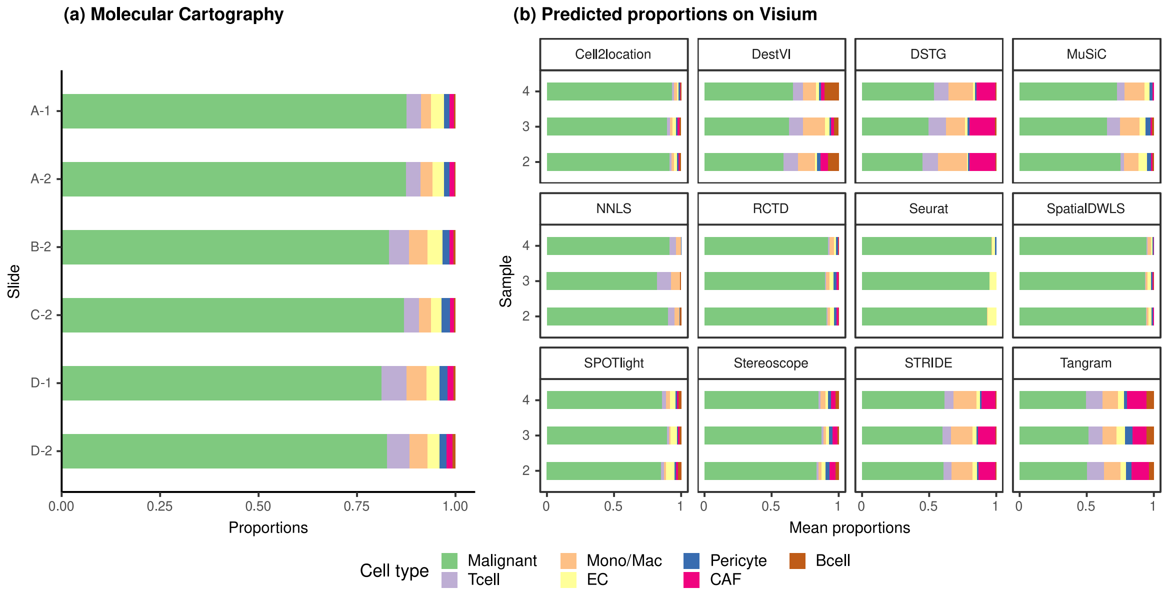


**Figure S16. (a)** Different melanoma sections profiled by Molecular Cartography have consistent cell type proportions, with an average inter- and intra-sample JSD of around 0.003. Sections with the same number belong to the same sample, i.e., A-1 and D-1. **(b)** The average cell type abundances predicted by each deconvolution method are also consistent on the three melanoma Visium slides.
