## Supplementary Notes for "Spotless: a reproducible pipeline for benchmarking cell type deconvolution in spatial transcriptomics"

### Supplementary Note 1: Description and validation of the simulation procedure

To generate synthetic spots, *synthspot* by default samples 2-10 cells from input scRNA-seq data. The counts from these cells are then summed up per gene then downsampled using the *downsampleMatrix* function from *DropletUtils* [1] to be within the given mean and standard deviation (20,000 ± 5,000 counts by default). The uniqueness of *synthspot* lies in the variable cell type frequency priors between abundance patterns, which determine the probability that a cell type will be sampled during spot generation. The nine abundance patterns used in our benchmark are made up of three characteristics, 1) the uniformity of each region, the 2) distinctness of cell types within each region, and 3) whether or not there are missing, dominant, or rare cell types (**Figure S1**). **Supplementary Notes Figure 1** depicts the simulation process in detail. For simplicity purposes, we have excluded the

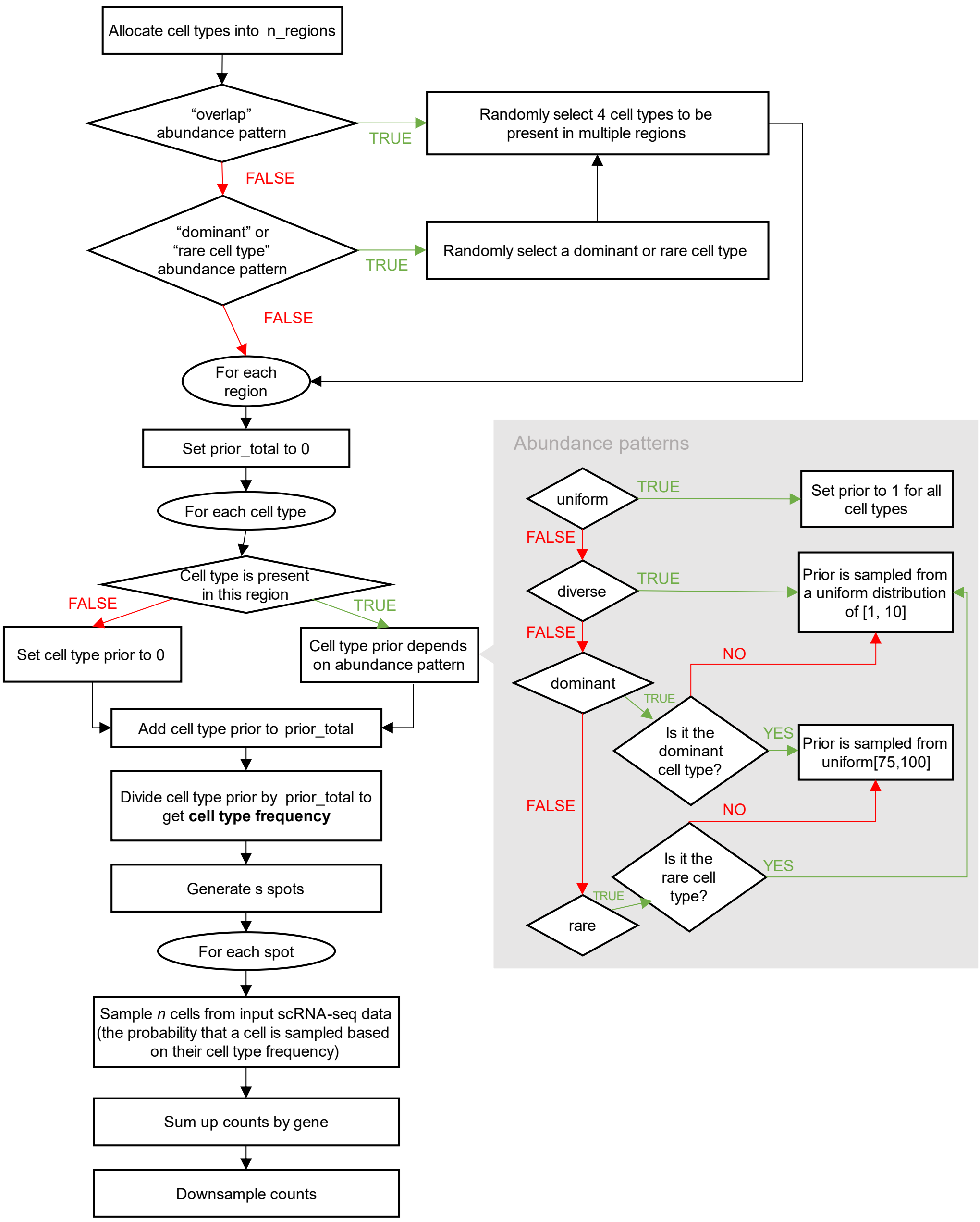


**Supplementary Notes Figure 1.** Schematic of the *synthspot* simulation algorithm

*partially dominant*, *regionally rare*, and *missing cell types* abundance patterns from this flowchart. For the *partially dominant* pattern, we would also randomly select a region the dominant cell type would be absent (cell type prior is then set to zero), and another region where it is as equally abundant as other cell types (prior is sampled from uniform distribution of [1,10]). Similarly, for the *regionally rare* pattern, we would select a select where the rare cell type will only be present in, and then the prior of the rare cell type would be set to zero for all other regions. For the *missing cell types* pattern, we would first randomly select four cell types to be removed. Note that it is also possible for synthspot to use the cell type composition of regionally annotated scRNA-seq data as frequency priors. In that case, it will create the number of regions equal to the scRNA-seq data and use cell type frequencies in the corresponding real region.

As an example, let us say we want to create a synthetic dataset containing two artificial regions which follow the *dominant cell type* pattern. Consider an input scRNA-seq dataset with 8 cell types called A, B, C, …, H. In this case, a dominant cell type is randomly selected which will be present in all regions (=H). Assume that region 1 contains cell types [A, B, C, E, H]. The abundance of H is sampled from a uniform distribution from 75 to 100, while the rest will be sampled from 1 to 10. Suppose we obtain the following priors: H = 80, A = 1, B = 2, C = 3, and E = 4. The summed abundance is 90, and the cell type frequencies are now: H = 0.9, A = 0.01, B = 0.02, C = 0.03 and E = 0.04. For all spots generated in region 1, these are the probabilities in which the cell types will be sampled.

We validated that our synthetic data and its abundance patterns sufficiently matches real Visium data in two ways, comparing the distributions with *countsimQC* [2] and using frequency priors based on real data. We compared synthspot with the algorithms to generate synthetic data used by cell2location, stereoscope, and SPOTlight. We used brain and kidney Visium datasets as the reference, and generated synthetic data using scRNA-seq data from the respective organs. These were the same scRNA-seq datasets used in our silver standard.

The counts per gene of each dataset seem to be representative of each algorithm's performance for other metrics, e.g., expression distribution, dispersion, mean-variance trend, and fraction of zeros (**Supplementary Notes Figure 2 & Supplementary Notes Figure 3**). For the brain dataset, synthspot has the most resemblance with real data, followed by SPOTlight (**Supplementary Notes Figure 2**). Since cell2location and stereoscope did not implement a downsampling step in their simulation, their synthetic brain datasets had overly abundant counts, a result of the plate-based scRNA-seq dataset (SMART-seq). SPOTlight downsamples each spot to have a total UMI count of 20,000, so the count distribution becomes uniform, unlike real data. For the kidney, all algorithms except stereoscope's were able to generate synthetic data that resembled real data (**Supplementary Notes Figure 3**). For most measures, cell2location's algorithm had the most resemblance with real data. Nonetheless, the robustness of synthspot towards sequencing technologies of the input dataset along with the flexibility in fine-tuning properties of the synthetic data makes it the preferred tool to aid in benchmarking. Finally, we verified that the distributions of the nine synthspot abundance patterns did not differ from one another visually or from the *real* abundance pattern, which uses real annotations as the frequency priors (**Supplementary Notes Figure 4**).

As a second verification, we used the regional annotation in the brain cortex dataset with the *real* abundance pattern to generate synthetic data with five brain regions (L1, L2/3, L4, L5, and L6), with each region having the same composition as the real layer. We then compared method performance between the *real* and *diverse overlap* patterns (**Supplementary Notes Figure 5**). Although the spot compositions between the patterns are different, method performances are similar, validating that artificial patterns can be used to evaluate model performance.

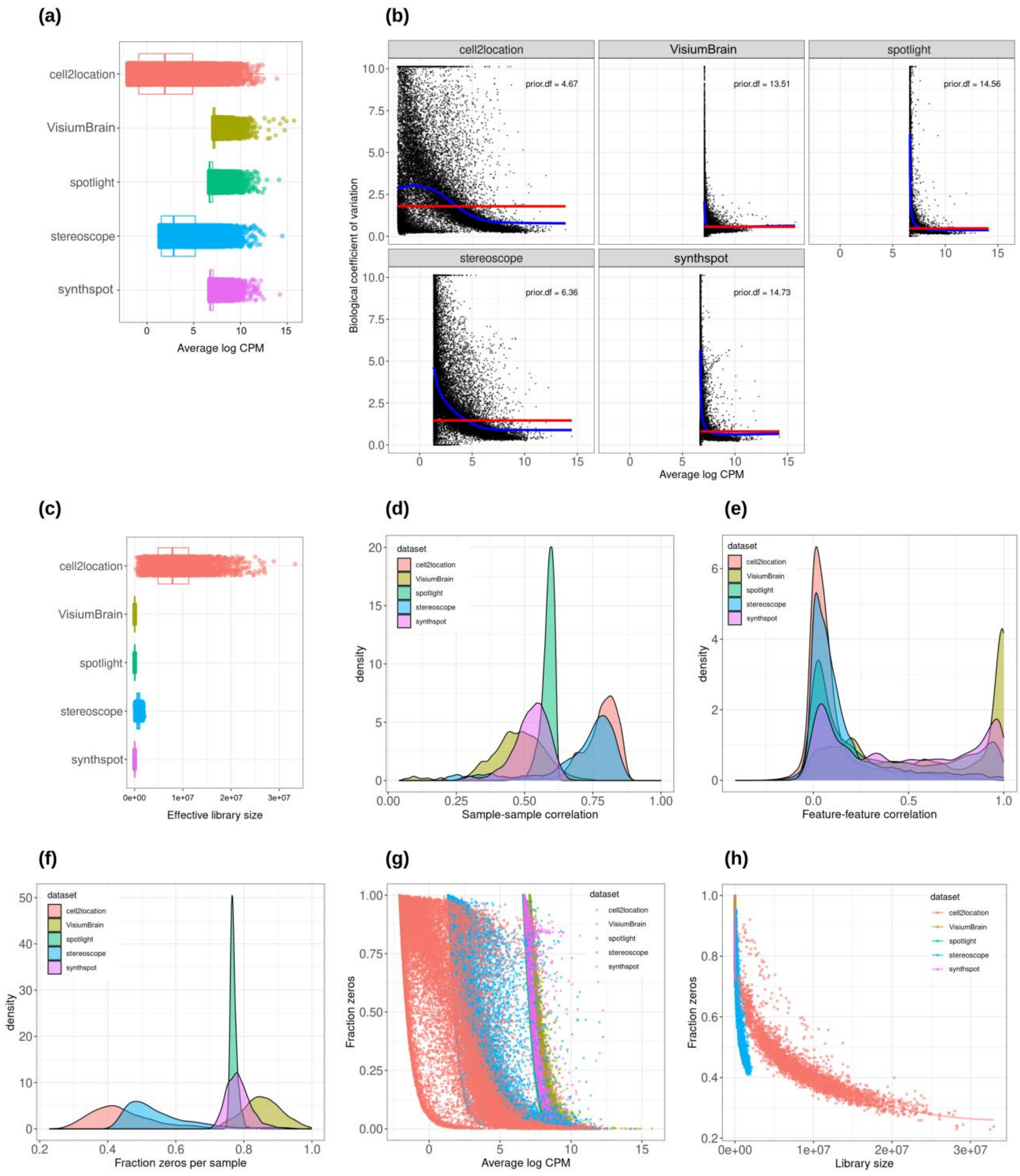


**Supplementary Notes Figure 2.** Plots comparing the characteristics of real Visium data from mouse brain and synthetic datasets generated from brain scRNA-seq data using different algorithms. **(a)** Average abundance values (log counts per million) per gene. **(b)** Association between average abundance and the dispersion. **(c)** Distribution of effective library sizes, or the total count per sample multiplied by the corresponding TMM normalization factor calculated by *edgeR*. **(d-e)** Distribution of pairwise Spearman correlation coefficients for 500 randomly selected spots **(d)** and genes **(e)**, calculated from log CPM values. Only non-constant genes are considered. **(f)** Distribution of the fraction of zeros observed per spot. **(g-h)** The association between fraction zeros and average gene abundance **(g)** and total counts per spot **(h)**.

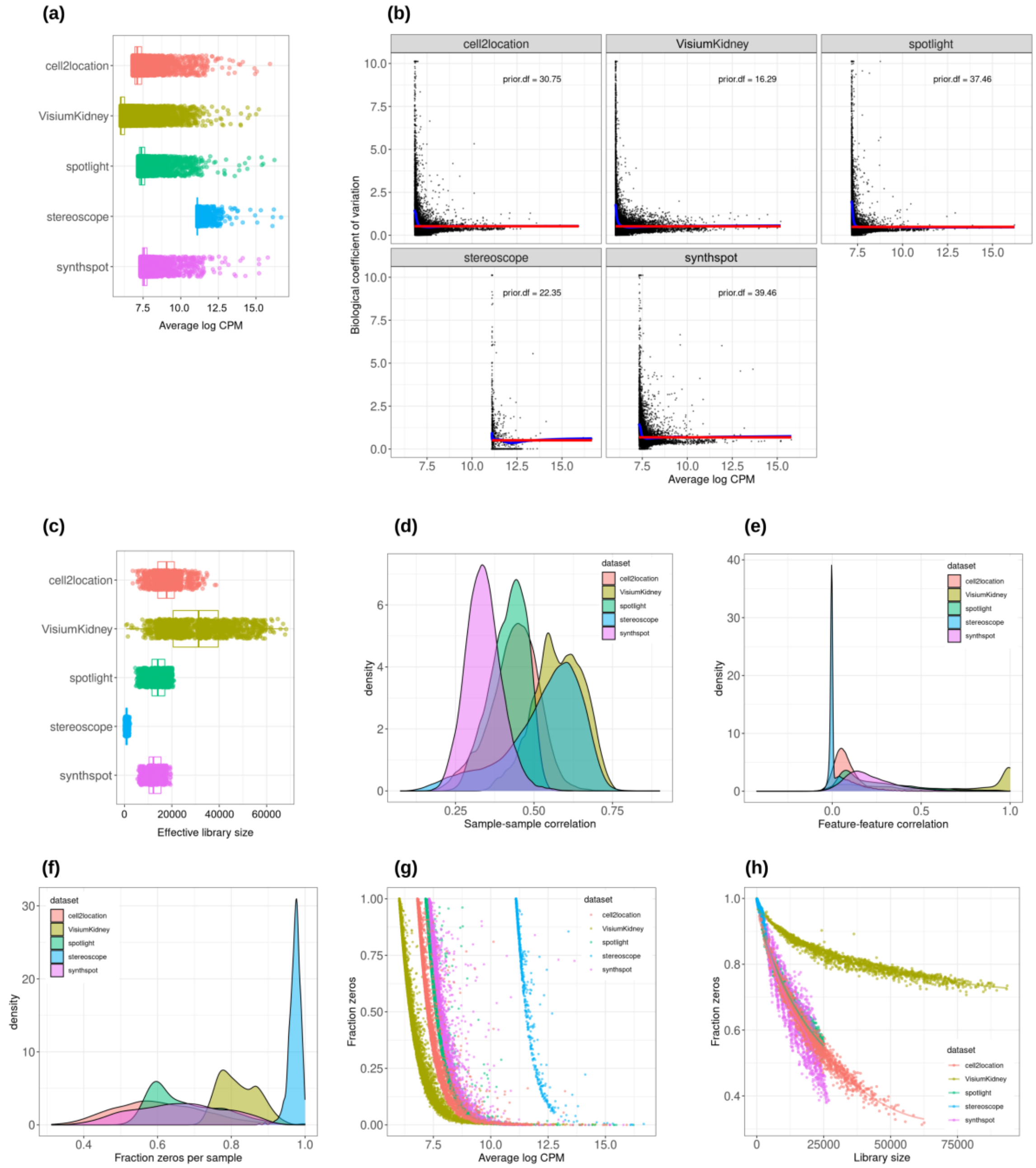


**Supplementary Notes Figure 3.** Plots comparing the characteristics of real Visium data from mouse kidney and synthetic datasets generated from kidney scRNA-seq data using different algorithms. **(a)** Average abundance values (log counts per million) per gene. **(b)** Association between average abundance and the dispersion. **(c)** Distribution of effective library sizes, or the total count per sample multiplied by the corresponding TMM normalization factor calculated by *edgeR*. **(d-e)** Distribution of pairwise Spearman correlation coefficients for 500 randomly selected spots **(d)** and genes **(e)**, calculated from log CPM values. Only non-constant genes are considered. **(f)** Distribution of the fraction of zeros observed per spot. **(g-h)** The association between fraction zeros and average gene abundance **(g)** and total counts per spot **(h)**.

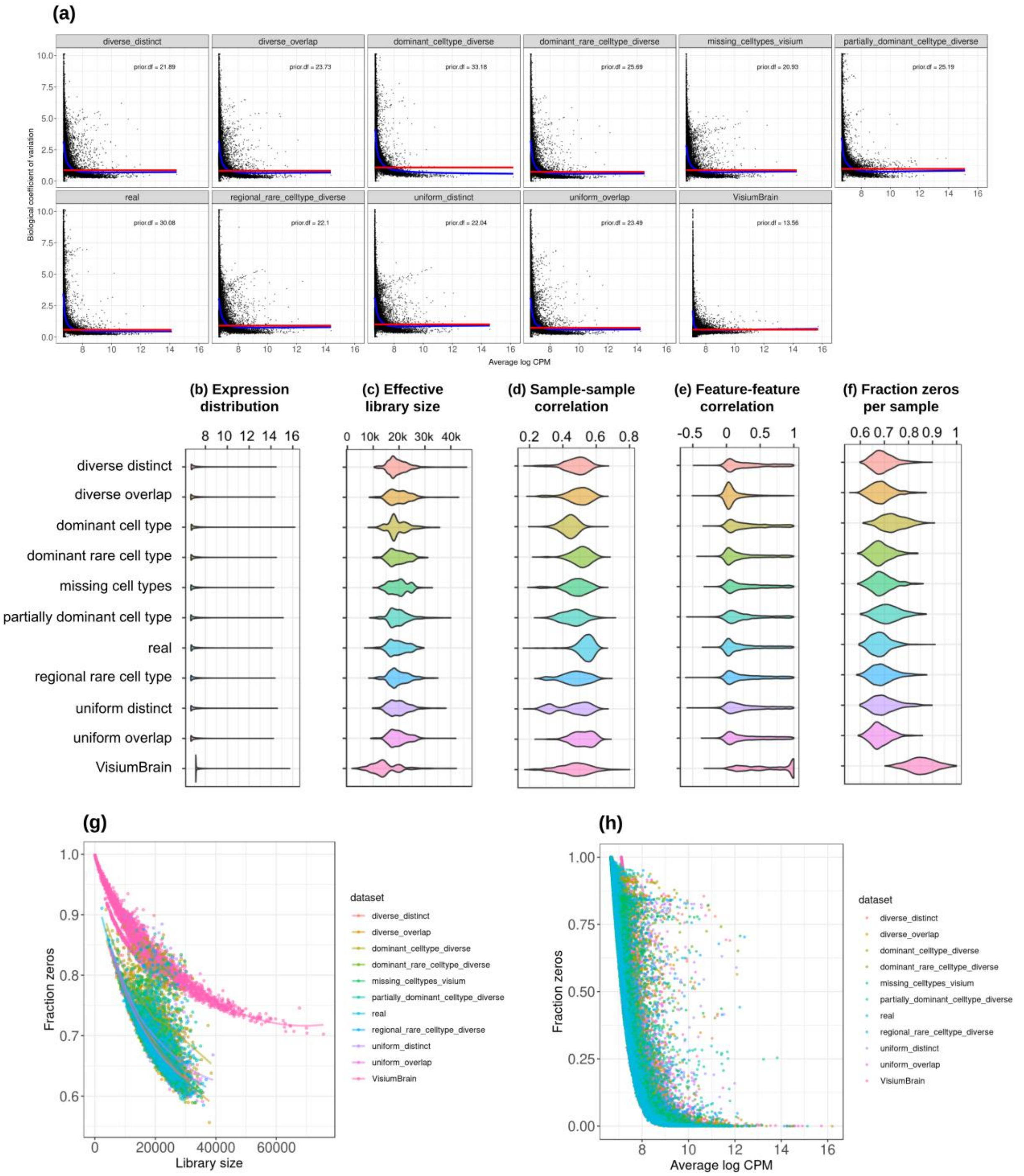


**Supplementary Notes Figure 4**. Plots comparing the characteristics of real Visium data from mouse brain and the eight synthetic abundance patterns from synthspot generated from brain scRNA-seq data. **(a)** Association between average abundance and the dispersion. **(b)** Average abundance values (log counts per million) per gene. **(c)** Distribution of effective library sizes, or the total count per sample multiplied by the corresponding TMM normalization factor calculated by *edgeR*. **(d-e)** Distribution of pairwise Spearman correlation coefficients for 500 randomly selected spots **(d)** and genes **(e)**, calculated from log CPM values. Only non-constant genes are considered. **(f)** Distribution of the fraction of zeros observed per spot. **(g-h)** The association between fraction zeros and average gene abundance **(g)** and total counts per spot **(h)**.

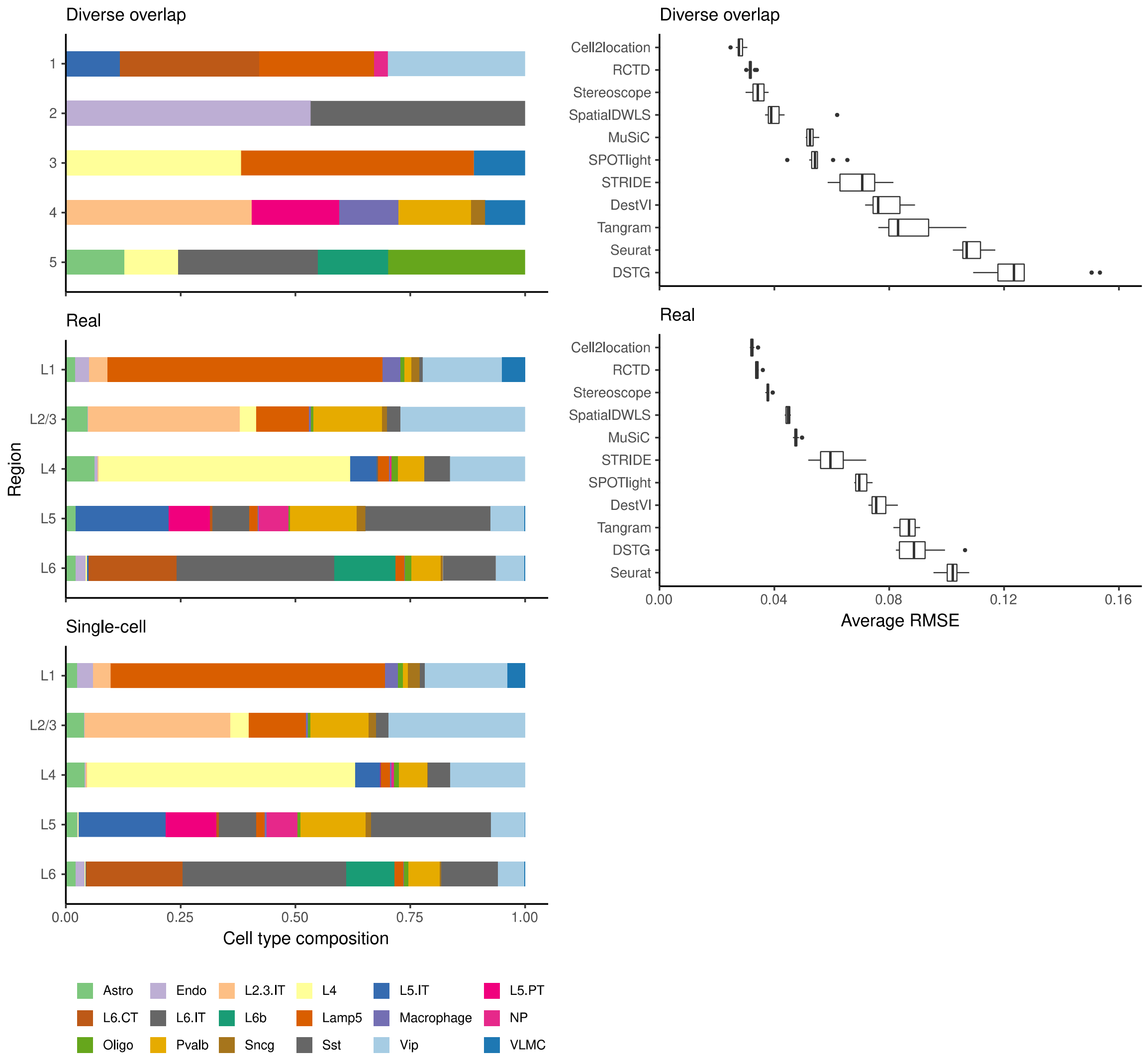


**Supplementary Notes Figure 5.** When using an *artificial* abundance pattern (*diverse overlap*) to create synthetic spatial data, method rankings remain almost identical as when using a *real* abundance pattern. The *real* pattern uses regional annotations from the scRNA-seq input to create regions with the same cell type frequencies.

### Supplementary Note 2: Issues with threshold-based classification metrics

In addition to the area under the precision-recall curve (AUPR), we initially included the (balanced) accuracy, specificity, recall (sensitivity), precision, and F1 score in the evaluation. These metrics evaluate the classification capabilities of each method, i.e., the correctness of cell type presence and absence prediction. Briefly, accuracy is the percentage of correctly classified cell types, specificity measures how many of the cell types predicted as absent are truly absent, sensitivity measures how well a method can detect a cell type within a spot, precision measures how many cell types predicted as present are truly present, and the F1 score integrates sensitivity and precision.

The first issue was that methods that use probabilistic models (e.g., cell2location, stereoscope, RCTD and DestVI) do not return proportions that are exactly zero but instead negligible values as low as $10^{-9}$. This made an unbiased evaluation difficult since a fixed threshold for cell type presence/absence must be selected to calculate classification metrics. In particular, different methods, datasets and abundance patterns have different thresholds for which the classification metrics are at a maximum.

The second issue stems from the class imbalance of our datasets, in which more cell types are absent than present in a spot (more negative than positive classes). In general, around 15% of the proportion matrix are positives classes, which made the specificity and precision particularly uninformative. This can be seen by how Seurat was the best performer on both specificity and precision despite having low sensitivity (**Supplementary Notes Figure 6**). By mostly predicting cell types to be absent in a spot, there are more false negatives (FNs), but specificity and precision do not take FNs into account. The balanced accuracy and F1 score were also unable to entirely correct for this class imbalance.

Given these two issues, we decided to use the precision-recall curve (PR) for evaluation instead and not include these five classification metrics (although we discuss its calculation below). The PR curve plots precision against recall at different thresholds, and the threshold to distinguish cell type absence/presence is varied from zero to one instead of fixing it at a certain value. Hence, the proportions are used as-is without rounding or binarization. It is also recommended for use with imbalanced datasets [3].

#### Calculation of threshold-based classification metrics

First, we rounded the predicted proportion matrices to two decimal points, so a proportion of 0.005 and under was rounded to zero. We calculated the micro-average of each classification metric. This is a global metric where the contributions of all classes are considered. We essentially treat the proportion matrix as in a binary classification problem and go through each element individually. As an example, the micro-precision is calculated as

$$Precision_{micro} = \frac{\sum_z^Z TP_z}{\sum_z^Z TP_z + \sum_z^Z FP_z}$$

where TP stands for true positive and FP for false positive. This is in contrast to the macro-average, where the metric is computed independently for each cell type using a one-vs-all approach, and the average is taken across all cell types ($Precision_{macro} = \frac{1}{Z}\sum_z^Z Precision_z$).

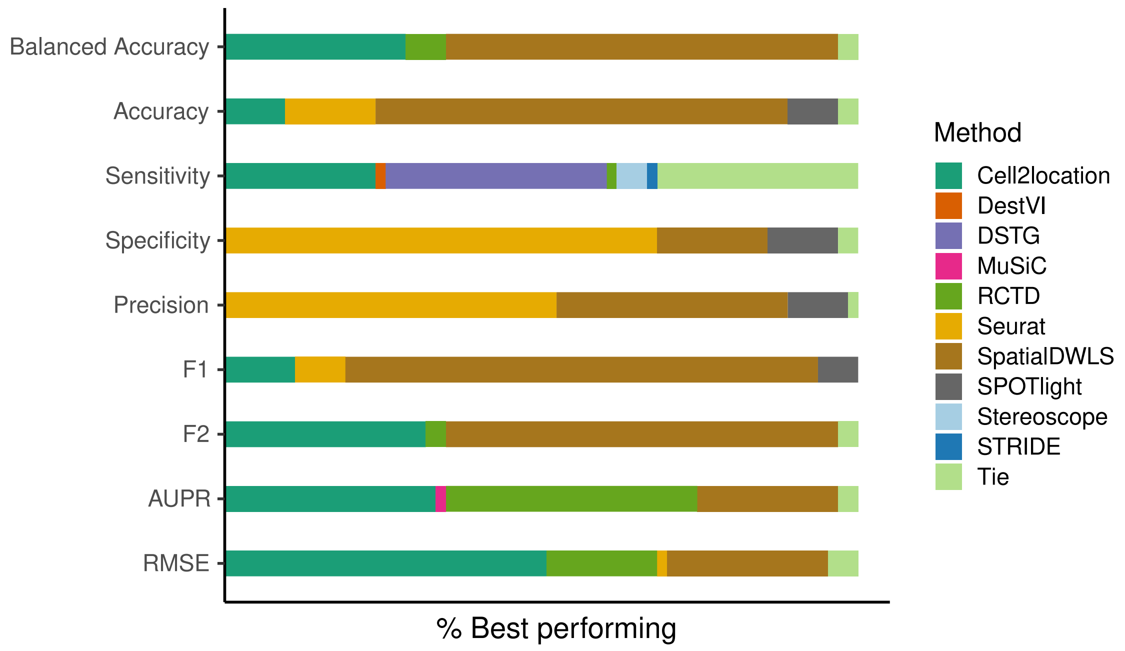


**Supplementary Notes Figure 6.** The relative frequency in which a method performs best in the silver standard, based on the best median value across ten replicates for that combination. Tie means that two or more methods score the same up to the third decimal point. RMSE: root-mean-square error; PRC AUC: area under the precision-recall curve.

### Supplementary Note 3: Method execution and parameter choice

In this supplementary note, we briefly describe each method, how we ran them, and the parameters that we evaluated. As most methods contain adjustable parameters that can affect its performance, we tested a range of parameter options to ensure optimal performance for each method. For reproducibility, users can find the exact parameters we have used for each analysis under the "conf/" folder in our Github repository.

*cell2location (v0.06a)*. Cell2location models the transcripts with a negative binomial distribution. It uses variational inference to estimate all the parameters. One advantage of cell2location is that users can shape model priors by providing hyperparameters that correspond to their prior knowledge. We used default priors from the tutorial of cell2location version 0.3. We filtered genes from the reference and spatial datasets as suggested by the tutorial. Model fitting was performed with cross-validation stratified by cell type annotation (*stratify_cv*). Sample information was not given to the model. The number of training iterations was set to 30,000 (*n_iter*).

In a newer tutorial, the default value of *detection_alpha* hyperparameter changed from 200 to 20. We compared the old and new values on the brain cortex seqFISH+ dataset (gold standard) and the kidney dataset (silver standard) and did not find a difference in performance (**Supplementary Notes Figure 7a**). Hence, we kept *detection_alpha=200* for our benchmark. We also varied the number of cells per location from 10-50 cells in the gold standard but also did not find a noticeable change in performance (**b**).

*DestVI (v0.16.0).* DestVI uses latent variable models (i.e., variational autoencoders) for the single-cell and spatial data. Unlike other methods, it also models a continuous estimate of cell state for every cell type in every spot. With the default of 2,500 epochs, training did not converge for many of the silver standard datasets. We followed the author's recommendations and increased to 5,000 training epochs and reduce the minibatch size (*batch_size*) to 64, which improved performance for all silver standard datasets (**Supplementary Notes Figure 8**). For the gold standards, we used 2,500 training epochs and *batch_size*=4.

*DSTG (v0.0.1).* DSTG uses a graph convolutional neural network to learn the composition of real spots from simulated spots. It performs a joint dimensionality reduction (canonical correlation analysis, CCA) and identifies the mutual nearest neighbors between the real and simulated spots. As many of the parameters were hardcoded, e.g., 200 nearest neighbors and 30 canonical vectors, the algorithm did not run on our gold standards where there were only nine spots per FOV. We adjusted the source code to change *k_filter*, *k*, *num_cc*, and *dims* to be equal to the number of spots in such cases, otherwise the default parameters were used.

*MuSiC (v0.2.0).* MuSiC is a bulk deconvolution method developed to handle multiple scRNA-seq references from multiple subjects. It employs weighted non-negative least squares (NNLS) regression. In case there are scRNA-seq reference datasets from multiple samples, the between-subject variance is used as weights for each gene. Genes with consistent expression among subjects are considered more informative and will be given higher weights during regression. We ran MuSiC without pre-grouping of cell types. We did not provide subject information to the model but instead considered each cell as a separate subject. This is not all our silver standards contained sample information, and we observed that the performance remained the same or worse when the sample information was given (**Supplementary Notes Figure 8a**). We also experimented with creating "pseudosamples" for datasets without sample information, where each cell was randomly assigned to 1 out of 3 artificial samples. This returned a worse performance than when single cells were used as samples (**b**).

*RCTD (v1.2.0)*. RCTD models the transcripts as being Poisson distributed and uses maximum likelihood estimation to infer cell type proportions. We ran RCTD with *doublet_mode="full"*, indicating that many cell types per spot were to be expected.

*Seurat Integration (v4.1.0).* Using joint dimensionality reduction between the scRNA-seq (reference) and spatial (query) data, we defined compatible reference-query pairs and used them as anchors for the label transfer procedure. We obtain a probability for a cell type being in the spot and use this as proxy for the abundance. We compared two normalization methods (SCTransform and vst) and two projection methods (PCA and CCA) and found that using SCTransform with PCA gave the best results (**Supplementary Notes Figure 10**). As with DSTG, we changed the number of neighbors and vectors (*k.score*, *k.weight*, *dims*) used in the gold standard as equal to the number of spots, otherwise the default was used.

*SpatialDWLS (v1.1.0).* SpatialDWLS performs cell-type enrichment analysis for each spot, then uses down-weighted least squares on marker genes. We followed the protocol described in Del Rossi et al. [4], where the *makeSignMatrixDWLS* function was used with top 100 marker genes, instead of following the online vignette where the *makeSignMatrixPAGE* function was used with all marker genes (**Supplementary Notes Figure 11**).

*SPOTlight (v0.1.7)*. SPOTlight is the only method based on non-negative matrix factorization (NMF) and NNLS. This method computes topics from the gene expression profile with NMF instead of using the expression values directly. NNLS is used with the topics to obtain the cell type proportions for each spot. We followed the vignette of version 0.1.5, as the recommended parameters have changed slightly between each version of SPOTlight. In this version, SPOTlight uses Seurat's *FindAllMarkers* functions to calculate marker genes between cell types, and the *logfc.threshold* (limit testing to genes which show at least X-fold difference) and *min.pct* (only test genes that are expressed in at least X% of cells) parameters have a huge impact on the resulting list of marker genes. Furthermore, the parameters *cl_n* (number of cell types to use) and *min_cont* (only keep cells that are at least X% present in a spot) within the deconvolution function itself affects the predictions. We tested three sets of parameters and in the end, ran *FindAllMarkers* with *only.pos=TRUE*, *logfc.threshold=1*, and *min.pct=0.9*, and the deconvolution with *cl_n=50* and *min_cont=0.09* (**Supplementary Notes Figure 12**). This was also the parameter set that gave the shortest runtime. We also normalized both the reference and spatial data with *SCTransform*.

*Stereoscope (v0.2.0)*. Stereoscope models the transcripts with a negative binomial distribution. It uses maximum likelihood estimation to infer the rate and overdispersion parameters from the scRNA-seq data and then uses maximum a posterior (MAP) estimation to infer cell type proportions. We ran stereoscope using all genes for the silver standard but only the top 5,000 most highly expressed genes for the gold standard, as that gave the best results for both cases. Although the authors have noted in their paper that choosing the 5,000 most expressed genes is sufficient, we saw that using all genes for the silver standards still gave slightly better performance (**Supplementary Notes Figure 13a**). We implemented an option to use the HVGs instead of top expressed genes, but this did not consistently result in better performance (**b**). Finally, we tested the *sub* parameter, which subsamples each cell type in the single-cell reference to at most X cells, but did not see any improvement in the kidney (silver standard) or liver dataset (**c**). We verified that both the training and test models have converged.

*STRIDE (v0.0.2).* STRIDE trains a topic model from scRNA-seq data, then applies this model to the spatial data. We ran it with *--normalize* and we let STRIDE automatically select the optimal topic number for each dataset. We did not find a pattern when comparing results from raw and normalized counts (**Supplementary Notes Figure 14**).

*Tangram (v1.0.3).* Tangram learns a spatial alignment of scRNA-seq from the spatial data via nonconvex optimization. Like Seurat integration, Tangram returns the probabilistic counts for each cell type in each cell voxel. We tested the three mapping modes: *cells* (maps single cells to spots), *clusters* (maps average of cells per cell type instead of single cells), and *constrained* (constrain the number of mapped single cell profiles). Although the *constrained* mode was recommended for deconvolution, we found that using the *clusters* resulted in the best performance (**Supplementary Notes Figure 15**). When running the *constrained* mode, we also provided the ground truth number of cells per spot and total number of cells in the *density_prior* and *target_counts* parameters, respectively. Increasing the training epochs did not have an effect on the performance, and we verified that the models have converged. The final parameters we used were *map_cells_to_space* with *mode="clusters"* and *density_prior="rna_count_based"*. We used the top 100 marker genes for each cell type.

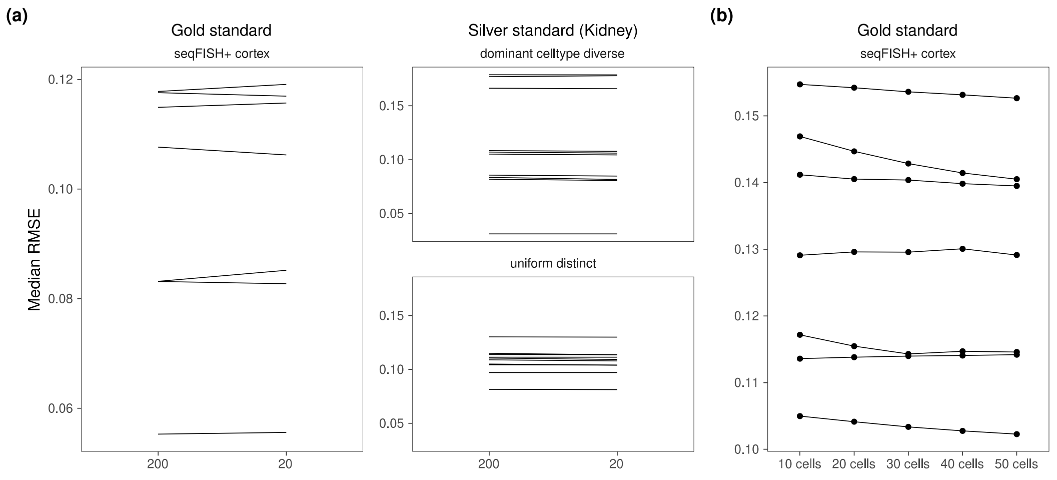


**Supplementary Notes Figure 7.** Changing hyperparameters in the cell2location model. There is almost no performance difference when changing the (a) detection alpha and (b) number of cells per spot.

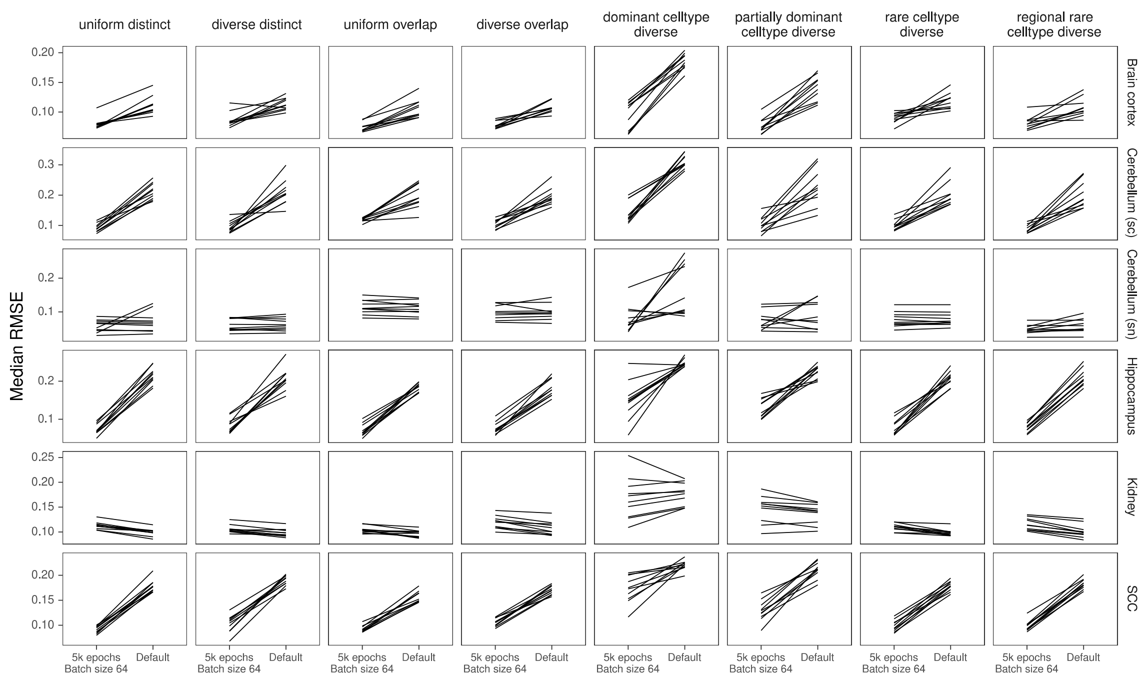


**Supplementary Notes Figure 8.** Compared to the default parameters (2500 epochs), DestVI has better performance with 5000 training epochs and batch size of 64.

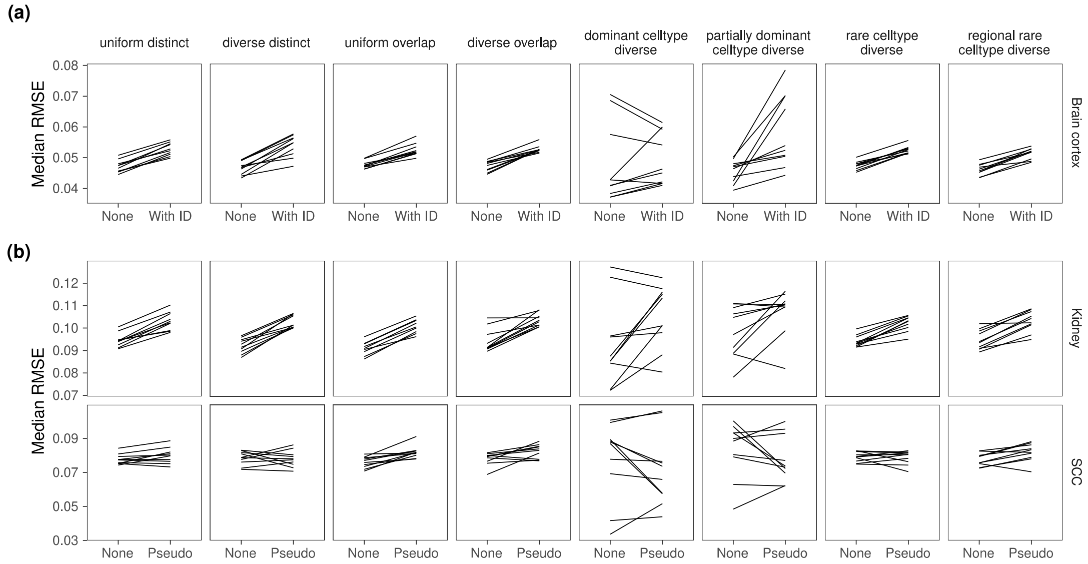


**Supplementary Notes Figure 9.** Comparing MuSiC performance when the model was given sample information. MuSiC seems to perform best when single cells were used as samples ("None"), as compared to (a) when the real sample information was given and (b) when pseudosamples were created.

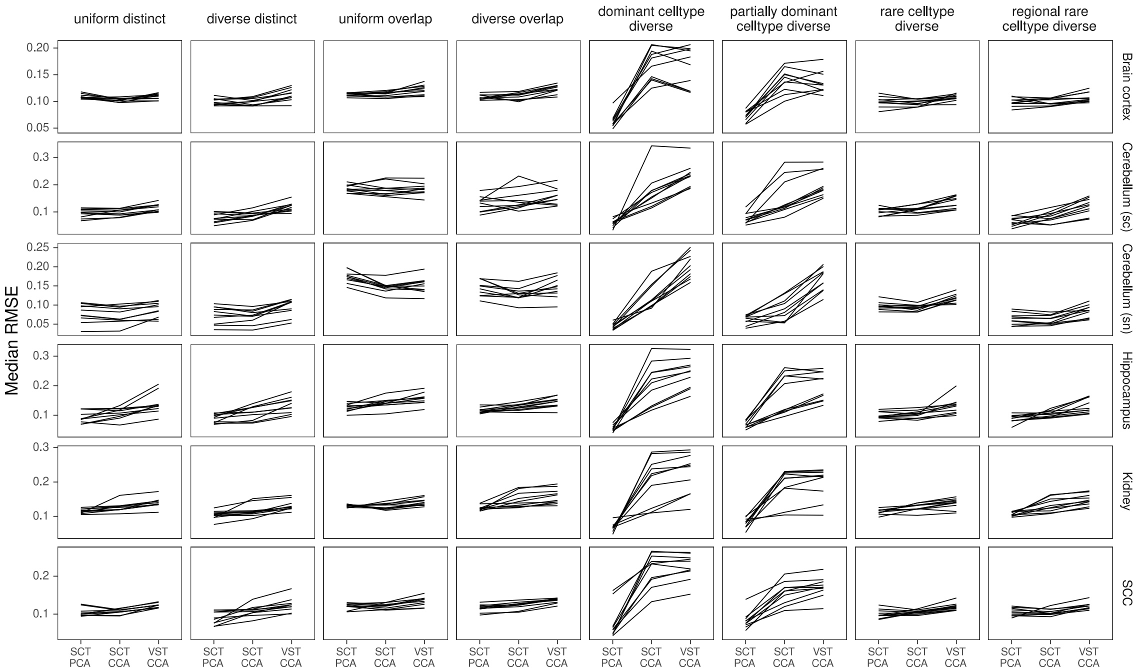


**Supplementary Notes Figure 10.** Seurat has the best performance when the data was transformed using SCTransform and the dimensionality reduction method is PCA. CCA = canonical correlation analysis; PCA = principal component analysis; VST = variance stabilizing transformation.

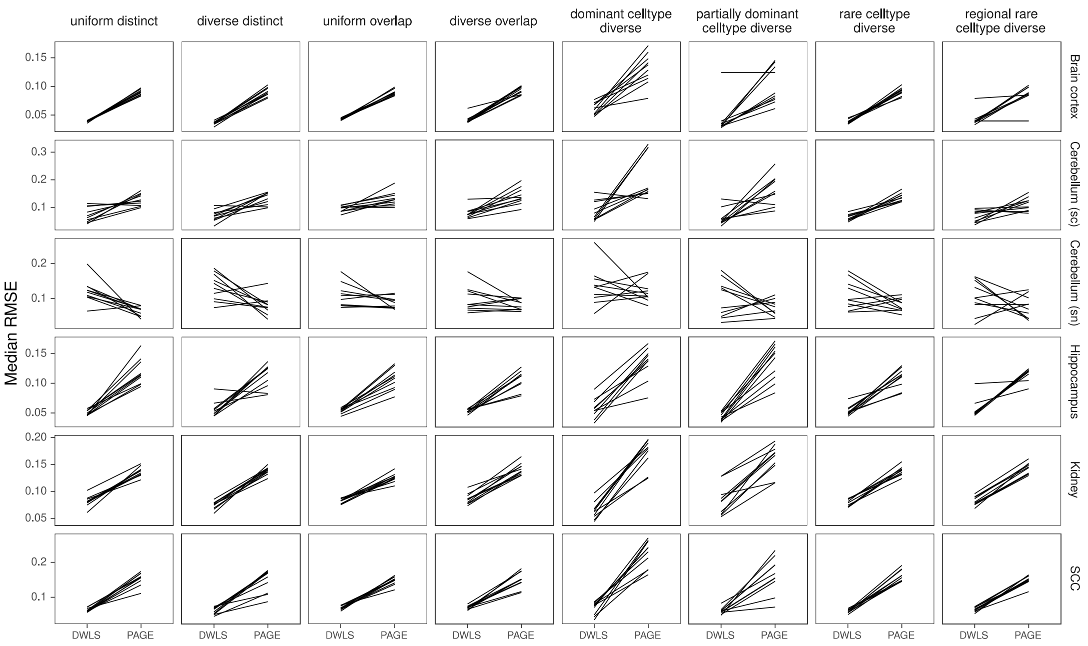


**Supplementary Notes Figure 11.** SpatialDWLS has better performance when the *makeSignMatrixDWLS* was used to create the signature matrix (as described in the Current Protocols paper), instead of the *makeSignMatrixPAGE* function (described in the online vignette).

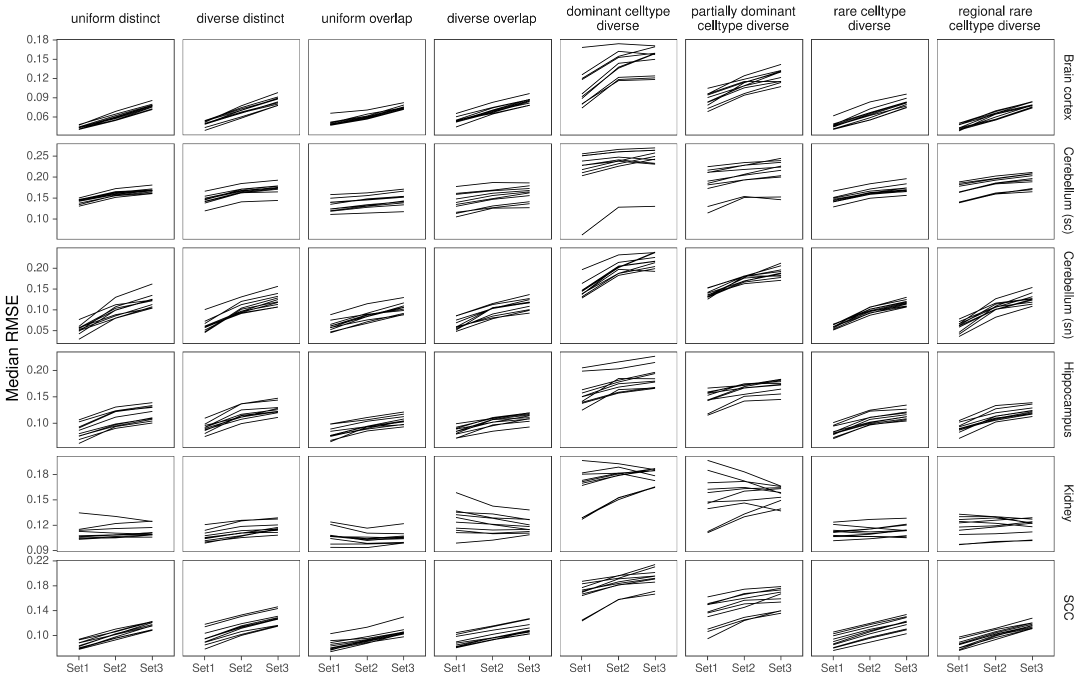


| | **logfc.threshold** | **min.pct** | **cl_n** | **min_cont** |
|---|---|---|---|---|
| *Set1* | 1 | 0.9 | 50 | 0.03 |
| *Set2* | 0.25 | 0.1 | 100 | 0 |
| *Set3* | 1 | 0.9 | 50 | 0.09 |

**Supplementary Notes Figure 12.** Three sets of parameters tested for SPOTlight. We used Set1 parameters in our benchmark.

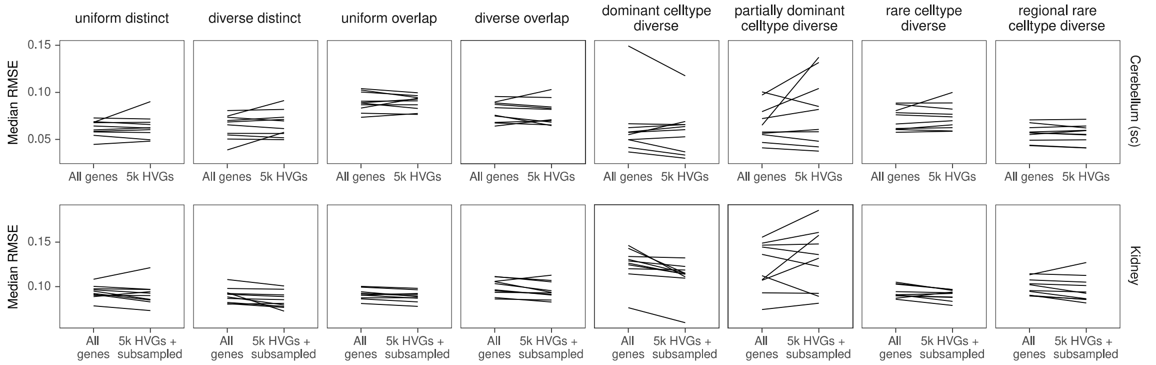


**Supplementary Notes Figure 13.** There is no consistent performance difference between using all genes of stereoscope and using the 5000 HVGs (with or without subsampling the scRNA-seq reference).

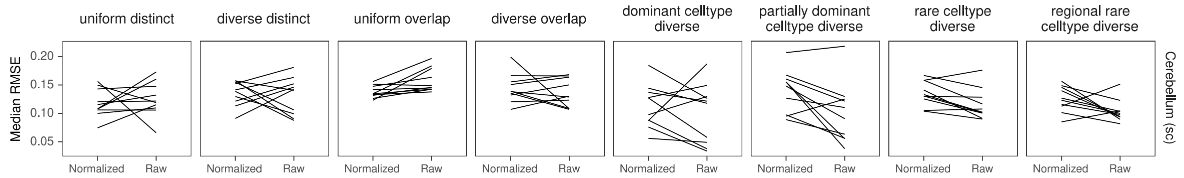


**Supplementary Notes Figure 14.** For STRIDE, there is no consistent performance difference between normalizing or using the raw counts.

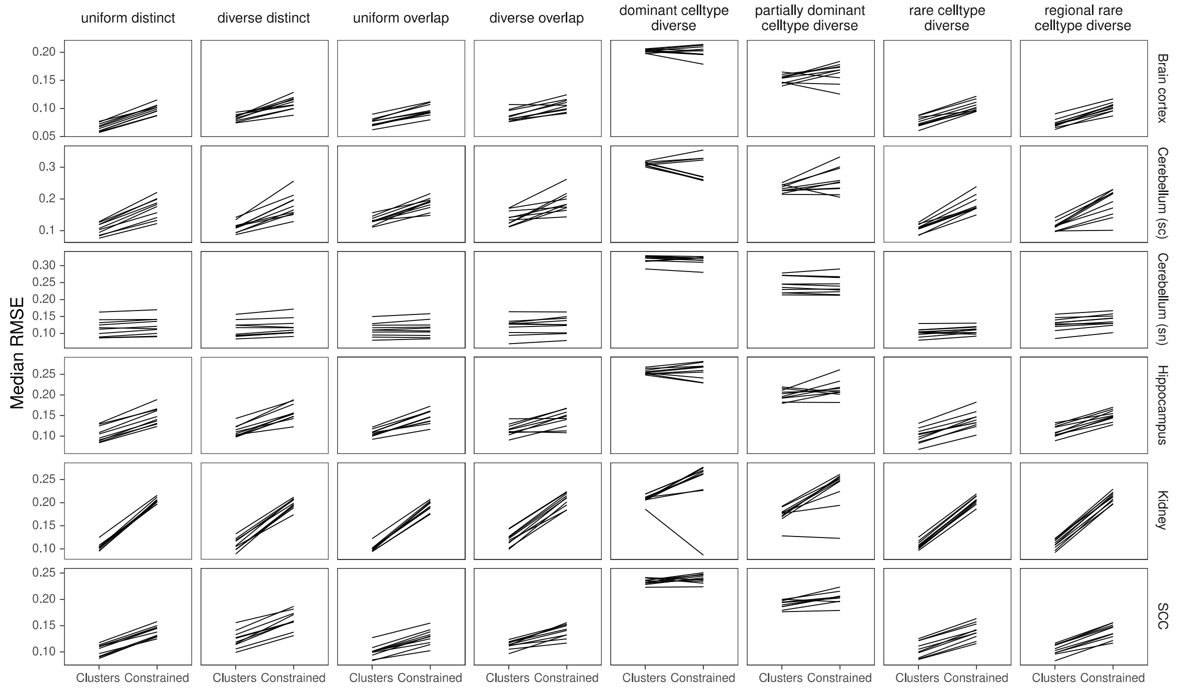


**Supplementary Notes Figure 15.** Although the *constrained* mapping mode was recommended in the Tangram vignette, we found that the *clusters* mode achieve better performance.
